# Distinct lifespan trajectories of cortical intrinsic timescales and neural oscillations

**DOI:** 10.64898/2026.11.04.764097

**Authors:** Xiaobo Liu, Brendan Harris, Runhao Lu, Sanwang Wang, Qiuxuan Yu, Mingyue Zhang, Ruifang Cui, Xinyu Liang, Tengteng Fan

**Affiliations:** McConnell Brain Imaging Centre, Montreal Neurological Institute, McGill University, Montreal, Canada; School of Physics, The University of Sydney, Sydney, NSW, Australia; Peking University Sixth Hospital, Peking University Institute of Mental Health, NHC Key Laboratory of Mental Health (Peking University), National Clinical Research Center for Mental Disorders (Peking University Sixth Hospital), Beijing, China; School of Information Engineering, Huzhou University, Huzhou, Zhejiang, China; Institute of Science and Technology for Brain-inspired Intelligence, Fudan University, Shanghai, China

**Keywords:** Cortical intrinsic timescales, Brain development and ageing, Magnetoencephalography, Corticothalamic dynamics, Neural oscillations, Computational modelling

## Abstract

The brain integrates information over time through the persistence of neural activity, yet how this persistence changes across the human lifespan and the mechanisms underlying these changes remain unclear. Using resting-state magnetoencephalography from three independent cohorts (n = 1,156; ages 6–88 years), we systematically examined lifespan changes in neural temporal dynamics, focusing on cortical intrinsic timescales and alpha oscillatory frequency. We found that cortical intrinsic timescales shortened from approximately 29 ms in childhood to 9 ms in late adulthood, with the steepest decline occurring before age 30. This compression was reproducible across cohorts and observed in all cortical regions, while the sensory-to-association hierarchy of timescales remained preserved throughout life. By contrast, alpha frequency followed a distinct trajectory, peaking in early adulthood before declining after midlife. Individualized corticothalamic models linked timescale compression to stronger intrathalamic inhibition, whereas increased conduction delays partly accounted for alpha slowing. Further network simulations indicated that intrinsic timescales primarily reflect collective network dynamics rather than single-neuron kinetics. Together, these findings reveal distinct lifespan trajectories of neural persistence and oscillatory rhythms, implicating collective corticothalamic dynamics in the temporal organization of the human brain across development and ageing.

## Introduction

Flexible behaviour requires the brain to combine what is happening now with what happened a moment ago. Speech comprehension, motor sequencing and working memory all depend on neural populations that hold information for tens to hundreds of milliseconds. How long cortical activity retains its own past—its intrinsic timescale—therefore sets the window over which information can be integrated, and changes in this window across life could reshape how the brain processes time. Intrinsic timescales are read from the decay of a population’s autocorrelation or, equivalently, from the ‘knee’ of the aperiodic component of its power spectrum^1,2^. Previous work established that timescales lengthen from sensory to association cortex in monkeys and humans^1,2,3^, that they are shaped by cortical microarchitecture, and that they are shorter in a small sample of older adults^2,4^. A recent synthesis^3^ emphasises that a measured timescale can arise from single-neuron properties or from recurrent network interactions, and that these two origins make different predictions. These findings characterise where timescales are long and how they could be generated; they do not tell us when across the lifespan the cortical integration window changes, or whether this change is the same process as the well-known slowing of brain rhythms.

This distinction matters because ‘speed’ is not one property of a neural circuit. A circuit’s rhythms set the pace at which it oscillates; its intrinsic timescale sets how long it remembers a perturbation^4,5^. The resting alpha rhythm slows from mid-life onwards and is widely used as an index of neural slowing^4^. If the alpha rhythm and the intrinsic timescale are two readouts of a single ageing process, they should change together across life and across individuals. If instead they are two clocks, they could change on different schedules and depend on different parts of the circuit that generates resting dynamics. Whether such a dissociation exists, and what organises it, remains unresolved.

Ageing and development provide a natural experiment for this question. Between childhood and old age, synaptic kinetics, inhibition, myelination and thalamocortical connectivity all change. If intrinsic timescales are a fundamental property of cortical organisation, they should reorganise systematically with age, and the pattern of this reorganisation—its timing, its spatial distribution and its relation to rhythms—should constrain which circuit mechanisms are plausible.

Candidate mechanisms fall into two classes. Timescales could be set within cortex, by synaptic kinetics and by the strength of local recurrent excitation and inhibition^3,5,6^. Alternatively, they could be set by loops that leave cortex. Cortical populations are embedded in reciprocal loops with thalamic relay nuclei, which are in turn inhibited by the thalamic reticular nucleus, and relay and reticular populations form an intrathalamic loop of their own^7–9^. These loops shape the rhythms of the waking and sleeping brain^7^ and sustain persistent cortical activity during working memory^10–12^, the kind of persistence that an intrinsic timescale measures. The thalamus also changes across life: its volume declines with age^13,14^. The two classes make different predictions. A change in synaptic kinetics or local gain acts on the cortical field alone. A delayed corticothalamic loop, in contrast, sets the frequency of the alpha resonance, and an intrathalamic feedback loop adds damping that cortex alone does not have.

Neural-field theory provides such a model. The corticothalamic model of Robinson and colleagues^15,16^ describes the cortical field, the thalamic relay and reticular nuclei and the loops among them with physiologically interpretable parameters, including cortical, corticothalamic and intrathalamic loop gains, synaptodendritic rates and the corticothalamic delay and predicts the MEG or EEG power spectrum from them. It has been fitted to spectra across arousal states^17^ and to age-related changes in the EEG^18^. In this framework a stronger intrathalamic loop and a longer corticothalamic delay leave different signatures on the spectrum, on the knee and on the alpha peak respectively (Fig. 1c). Importantly, the model nests a cortex-only model: setting the thalamic loop gains to zero removes the thalamus. The need for a thalamic term can therefore be tested rather than assumed, by asking whether the nested cortical-only model accounts for the same spectra and the same age effects.

**Fig. 1.**
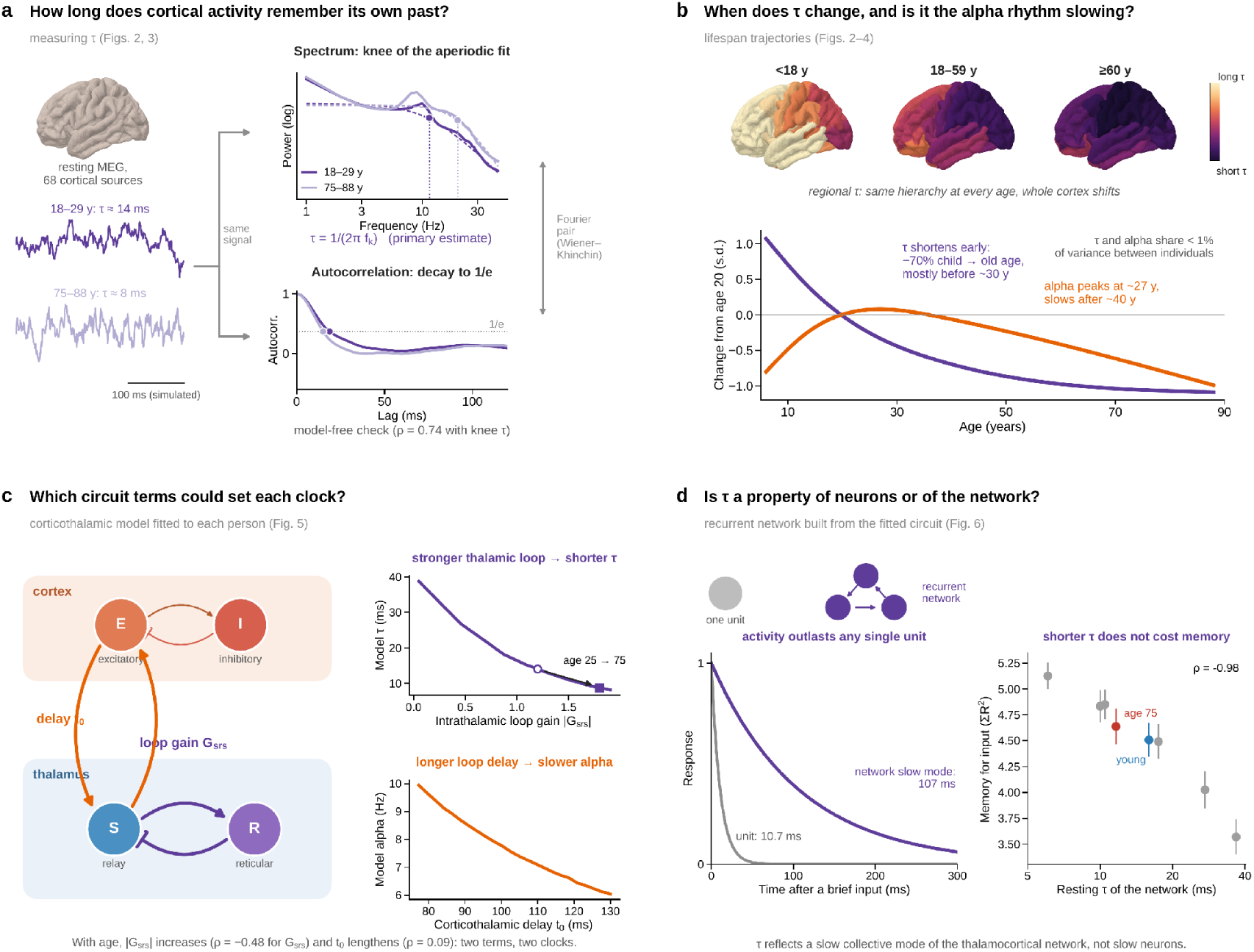
| Framework: intrinsic timescales, the alpha rhythm and the circuit terms that could set each. **a,** How long does cortical activity remember its own past? Resting-state MEG source activity (illustrated by simulated traces with the knee timescales of young, 18–29 years, and older, 75–88 years, Cam-CAN adults) has two equivalent read-outs, related by the Wiener–Khinchin theorem. Top, group-mean power spectra (solid) and aperiodic fits (dashed); the knee frequency *f*_k_ gives τ = 1/(2π*f*_k_), the primary estimate (13.8 and 7.9 ms for these group-mean spectra). Bottom, autocorrelation computed from the same spectra; its decay to 1/*e* is a check that does not assume a parametric knee model (18.7 and 15.0 ms; across participants, Spearman ρ = 0.74 with knee τ). **b,** When does τ change, and is it the same process as alpha slowing? Top, regional τ in youth (<18 years), adulthood (18–59 years) and older adulthood (≥60 years): the spatial hierarchy is present at every age while the whole cortex shifts to shorter τ. Bottom, cohort-adjusted trajectories of log τ and alpha peak frequency (change from age 20, s.d. units): τ shortens mostly before about 30 years, whereas alpha frequency peaks at about 27 years and slows after about 40 years. **c,** Which circuit terms could set each clock? A corticothalamic neural-field model (cortical excitatory, E, and inhibitory, I, populations; thalamic relay, S, and reticular, R, nuclei) is fitted to each participant’s spectrum. Right, model sweeps: a stronger intrathalamic loop gain |*G*_srs_| shortens model τ (open circle and filled square, fitted values at 25 and 75 years), and a longer corticothalamic delay *t*_0_ slows alpha frequency. **d,** Is τ a property of single units or of the network? A recurrent network built from the fitted circuit responds to a brief input with a slow collective mode (decay time ≈ 107 ms) that outlasts every single unit (≤ 10.7 ms in the young-adult network). Right, across eight network configurations a shorter resting τ goes with greater memory for input (mean ± s.d.; young-adult and age-75 networks highlighted).

Here we asked how cortical timescales change from childhood to old age using resting-state MEG source spectra from three independent cohorts (1,156 participants, 6–88 years). We first established the phenomenon and tested it against alternative explanations, including an autocorrelation-based estimate that does not assume a parametric knee model and 40 analytic specifications. We then asked where in cortex it occurs, whether it is the same process as alpha slowing, whether thalamic loops are needed to account for the spectra, which terms of the fitted corticothalamic model covary with the change, and using a recurrent network built from the fitted model, whether the timescale is a property of single units or of the network, and whether its compression must degrade temporal integration.

## Results

### Three cohorts, one spectral measure of the integration window

We analysed source-resolved resting-state magnetoencephalography (MEG) spectra from three independent cohorts comprising 1,156 participants aged 6–88 years: Cam-CAN (n = 606, 18–88 years, 299 females), OMEGA (n = 162, 18–74 years, 88 females) and PANS (n = 388, 6–85 years, 183 females; Supplementary Table 1). Power spectra were reconstructed for 68 Desikan–Killiany cortical regions, with frequency assignments and regional ordering verified across cohorts (Supplementary Fig. 1). Cortical intrinsic timescales (τ) were estimated from the knee frequency of the aperiodic spectral component over 2–45 Hz (Fig. 1a). Spectral parameterization showed a median fitting R² of 0.987 across 78,608 participant–region spectra, with the distribution of valid knee estimates and fitting quality summarized in Supplementary Fig. 2. Participant-level timescales were defined as the median across cortical regions with valid knee estimates. All 1,156 participants contributed to the primary analysis, and imposing a stricter criterion of at least 34 valid cortical regions yielded a consistent age association (Supplementary Table 2).

### Cortical timescales shorten by two-thirds across the lifespan, most steeply before about age 30

Between younger and older adults the cortex-averaged aperiodic spectrum kept relatively more power at higher frequencies—its low-frequency ‘shoulder’ moved rightwards (Fig. 2a). Expressed as a timescale, the median participant τ fell from 28.9 ms in children under 13 years to 15.9 ms at 18–24 years and 8.7 ms from age 75 (Fig. 2b,f). The association held within each cohort (Cam-CAN ρ = −0.43, OMEGA ρ = −0.33, PANS ρ = −0.64; all P < 2 × 10⁻⁵), with a within-cohort meta-analytic ρ = −0.50 (95% CI −0.54 to −0.45; Fig. 2c). Heterogeneity (I² = 93%) concerned magnitude, not direction: the effect was largest in PANS, the only cohort that includes children. Adjusting for sex and cohort, τ shortened by 11.7% per decade (95% CI 10.5–12.9%; partial η² = 0.21).

**Fig. 2.**
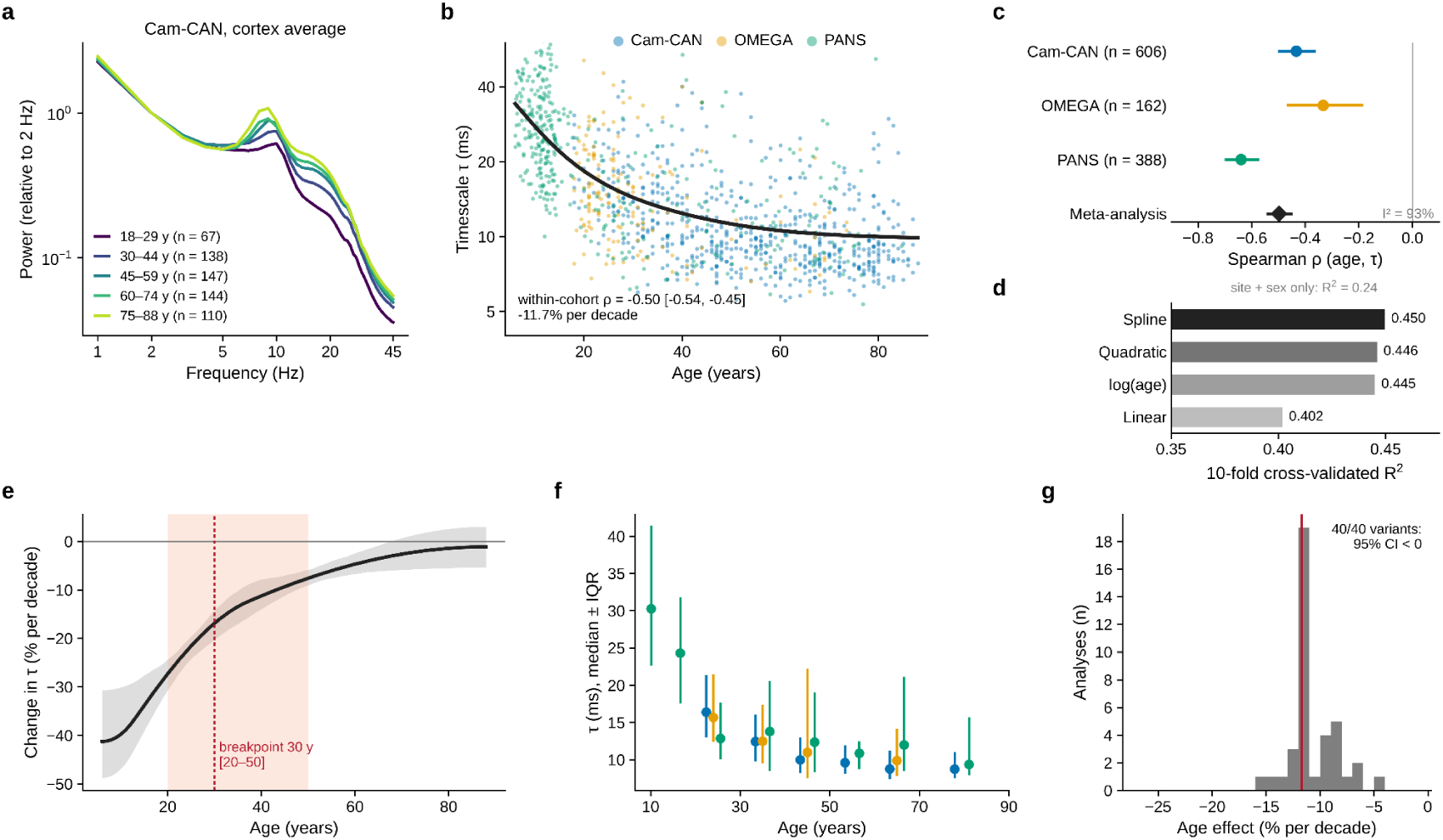
| Cortical intrinsic timescales shorten by about two-thirds across the lifespan, most steeply before about age 30. **a,** Cortex-averaged MEG power spectra for five age groups in Cam-CAN (18–29, 30–44, 45–59, 60–74 and 75–88 years), normalised to power at 2 Hz. **b,** Participant-level τ against age in three cohorts (Cam-CAN, *n* = 606; OMEGA, *n* = 162; PANS, *n* = 388; *N* = 1,156). Black curve, fitted non-linear trajectory adjusted for cohort and sex. The within-cohort meta-analytic Spearman ρ is −0.50 (95% CI −0.54 to −0.45), equivalent to an 11.7% reduction in τ per decade. **c,** Within-cohort Spearman correlations of age with τ (bars, 95% CI); the diamond is the fixed-effect meta-analytic estimate. All three cohorts are negative despite heterogeneity in magnitude (*I*² = 93%). **d,** Ten-fold cross-validated *R*² of four age models: spline 0.450, quadratic 0.446, logarithmic 0.445 and linear 0.402. **e,** Rate of change of τ (% per decade) from the spline model (black; grey band, bootstrap 95% CI). The dashed red line marks the breakpoint at 30 years from a two-segment regression (red band, 95% CI 20–50 years). **f,** Median τ (points) and interquartile range (bars) across age bins in each cohort. **g,** Age effect across 40 analytic specifications (the primary analysis and 39 variants; Supplementary Fig. 5); red line, primary analysis. All specifications are negative, with 95% confidence intervals that exclude zero.

The decline was not linear. A spline predicted held-out data better than a linear model (10-fold cross-validated R² = 0.450 versus 0.402; cohort and sex alone, R² = 0.24; Fig. 2d). A two-segment model placed the change in slope at 30 years (bootstrap 95% CI 20–50): τ shortened by 30% per decade before this point and by 6.4% per decade afterwards (both P < 10⁻¹¹; Fig. 2e). Most of the lifespan compression is therefore developmental and early-adult rather than a late-life decline. The breakpoint itself is imprecisely estimated (95% CI 20–50 years), so we interpret it as a transition around the third decade and not as a sharp boundary.

To determine whether timescale compression depended on spectral parameterization, we estimated the characteristic autocorrelation decay time from the same power spectra, without assuming a parametric knee model, using the Wiener–Khinchin relationship. Because the same spectra are used, this tests dependence on the parametric model, not on the recording. Autocorrelation-derived and knee-derived timescales were strongly associated across individuals (within-cohort ρ = 0.74). Autocorrelation timescales also shortened with age (ρ = −0.55; −4.5% per decade), with negative age associations in all 68 cortical regions and a regional age-effect map corresponding to that obtained from the spectral knee (ρ = 0.73; Supplementary Fig. 3). The negative association between age and intrinsic timescales was observed within all three cohorts (Supplementary Fig. 4; Supplementary Table 3a) and remained robust across alternative inclusion criteria, spectral fitting parameters, cohort adjustments and statistical models (Supplementary Fig. 5; Supplementary Table 2). Across the 40 analytic specifications reported in Supplementary Table 2 (the primary analysis and 39 variants), all estimated age effects were negative, with 95% confidence intervals excluding zero (range, −4.4% to −28.0% per decade). Although effect magnitudes varied across specifications, these analyses consistently supported the direction of lifespan-related timescale compression.

### The hierarchy sets where timescales are long at every life stage; the shortening is cortex-wide

Having established the phenomenon, we asked how it is organised across cortex. We mapped regional τ separately in youth (<18 years, n = 232), adulthood (18–59 years, n = 608) and older age (≥60 years, n = 316), and across all ages (Fig. 3a). The same spatial pattern was present at each stage (map similarity ρ = 0.90 between youth and adults, 0.76 between adults and older adults; Fig. 3b), while its overall level fell from a median regional τ of 36.1 ms in youth to 13.6 ms in adults and 10.0 ms in older adults. Timescales were shortest in superior parietal and occipital cortex and longest in orbitofrontal and temporal-pole cortex. The map followed the principal axis of functional connectivity from visual to transmodal cortex in youth (ρ = 0.53, P_spin_ = 0.008) and in adults (ρ = 0.57, P_spin_ = 0.003), replicating the invasive hierarchy of timescales, but this alignment was weaker and not significant in older adults (ρ = 0.37, P_spin_ = 0.23; Fig. 3c). Per-cohort maps were similar across cohorts (Supplementary Fig. 6).

**Fig. 3.**
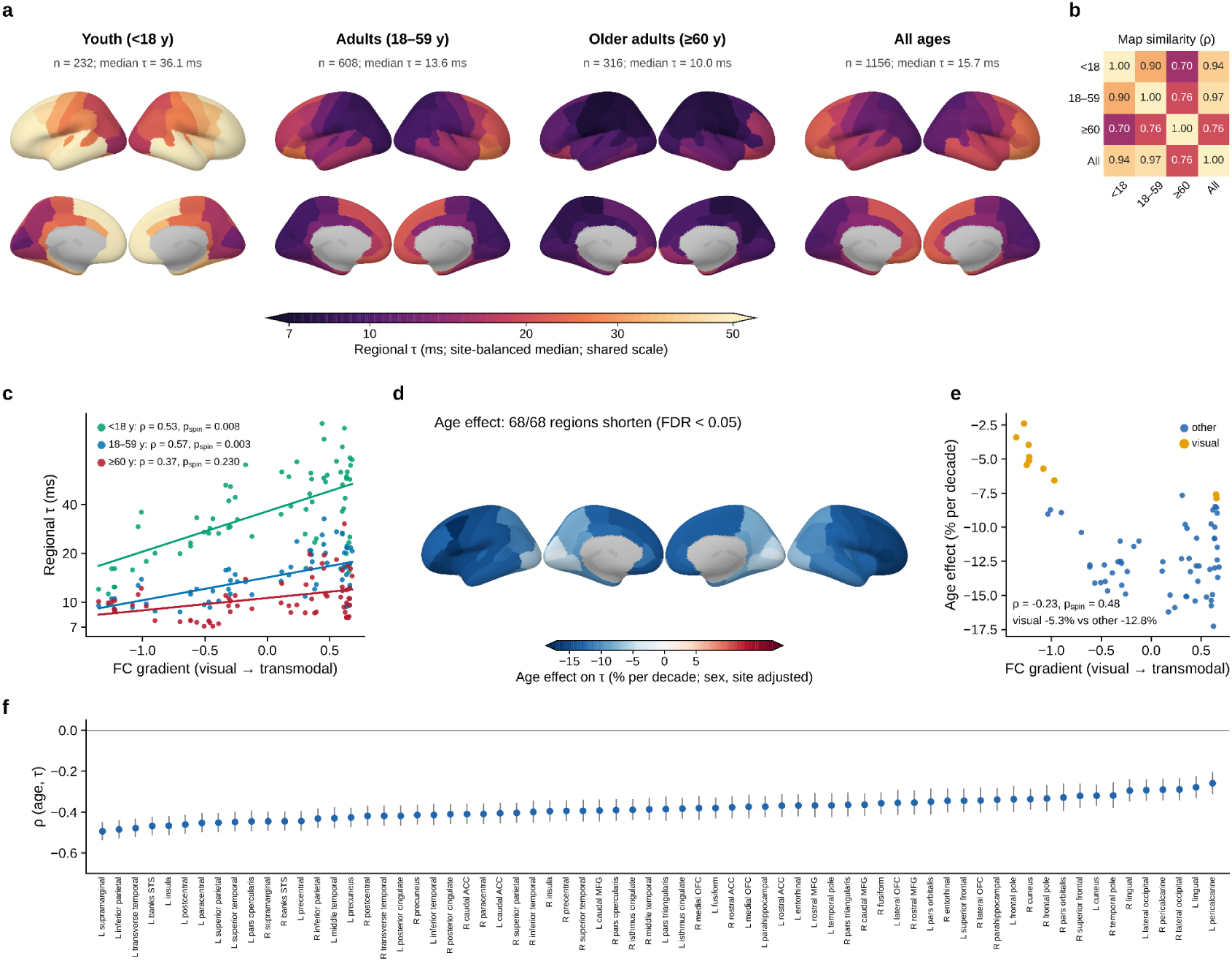
| The timescale hierarchy is preserved across life despite cortex-wide shortening. **a,** Regional τ across 68 Desikan–Killiany regions in youth (<18 years; *n* = 232), adults (18–59 years; *n* = 608), older adults (≥60 years; *n* = 316) and the full sample (*n* = 1,156): cohort-balanced medians on a shared logarithmic colour scale. Median regional τ falls from 36.1 ms in youth to 13.6 ms in adults and 10.0 ms in older adults. **b,** Spatial similarity between age-group maps (Spearman ρ): youth–adults 0.90, adults–older adults 0.76, youth–older adults 0.70. **c,** Regional τ against the principal functional-connectivity gradient (unimodal visual to transmodal), with linear fits per age group. The correspondence is significant in youth (ρ = 0.53, *P*_spin_ = 0.008) and adults (ρ = 0.57, *P*_spin_ = 0.003) but not in older adults (ρ = 0.37, *P*_spin_ = 0.230); differences between age groups were not significant under spin permutation (see text). **d,** Regional age effects on τ (% per decade, adjusted for sex and cohort); all 68 regions are negative after false-discovery-rate correction (FDR < 0.05). **e,** Regional age effects against gradient position (ρ = −0.23, *P*_spin_ = 0.48). Visual regions (orange) show smaller reductions (−5.3% per decade) than other regions (blue; −12.8% per decade). **f,** Region-specific within-cohort Spearman correlations of τ with age, combined by meta-analysis (points, ρ; bars, 95% CI; regions ordered by effect size).

The magnitude of age-related timescale compression was not systematically organized along the principal functional gradient. All 68 cortical regions exhibited negative age associations that survived false discovery rate correction (FDR < 0.05; Fig. 3d,f), with estimated reductions ranging from 2.4% per decade in the left pericalcarine cortex to 17.3% per decade in the left caudal middle frontal cortex. Regional age effects were not significantly associated with the functional gradient (ρ = −0.23, P_spin_ = 0.48; Fig. 3e). Nevertheless, early visual regions, including pericalcarine, cuneus, lingual and lateral occipital cortices, showed relatively smaller reductions than the remaining regions (median 5.3% versus 12.8% per decade). Thus, timescale compression was widespread across cortex, with spatial heterogeneity superimposed on a broadly preserved regional hierarchy. The spatial correspondence between intrinsic timescales and the principal functional gradient was numerically weaker in older adults (ρ = 0.37) than in adults (ρ = 0.57) or youth (ρ = 0.53). Spin-permutation tests of these differences were not significant (adults minus older adults, Δρ = 0.20, *P*_spin_ = 0.32; youth minus older adults, Δρ = 0.16, *P*_spin_ = 0.53), although participant-level bootstrap intervals excluded zero (95% CI 0.07 to 0.30 and 0.02 to 0.24). We therefore regard weaker alignment in later life as suggestive and not established.

### Intrinsic timescales and alpha frequency exhibit dissociable lifespan trajectories

We next asked whether age-related compression of intrinsic timescales reflected the same dimension of neural temporal dynamics as changes in alpha oscillatory frequency. The two measures followed contrasting lifespan trajectories (Fig. 4a,b; Supplementary Fig. 4; Supplementary Tables 3a and 3b). Intrinsic timescales shortened most rapidly during development and early adulthood, with within-cohort age correlations of ρ = −0.26 in youth (<18 years), −0.24 in early adulthood (18–39 years) and −0.17 after age 40. In contrast, alpha peak frequency increased during youth (ρ = 0.17), showed little age dependence between 18 and 39 years (ρ = −0.05), and declined after age 40 (ρ = −0.28). The fitted alpha trajectory reached its maximum at approximately 27 years, by which time intrinsic timescales had already declined substantially from childhood values.

**Fig. 4.**
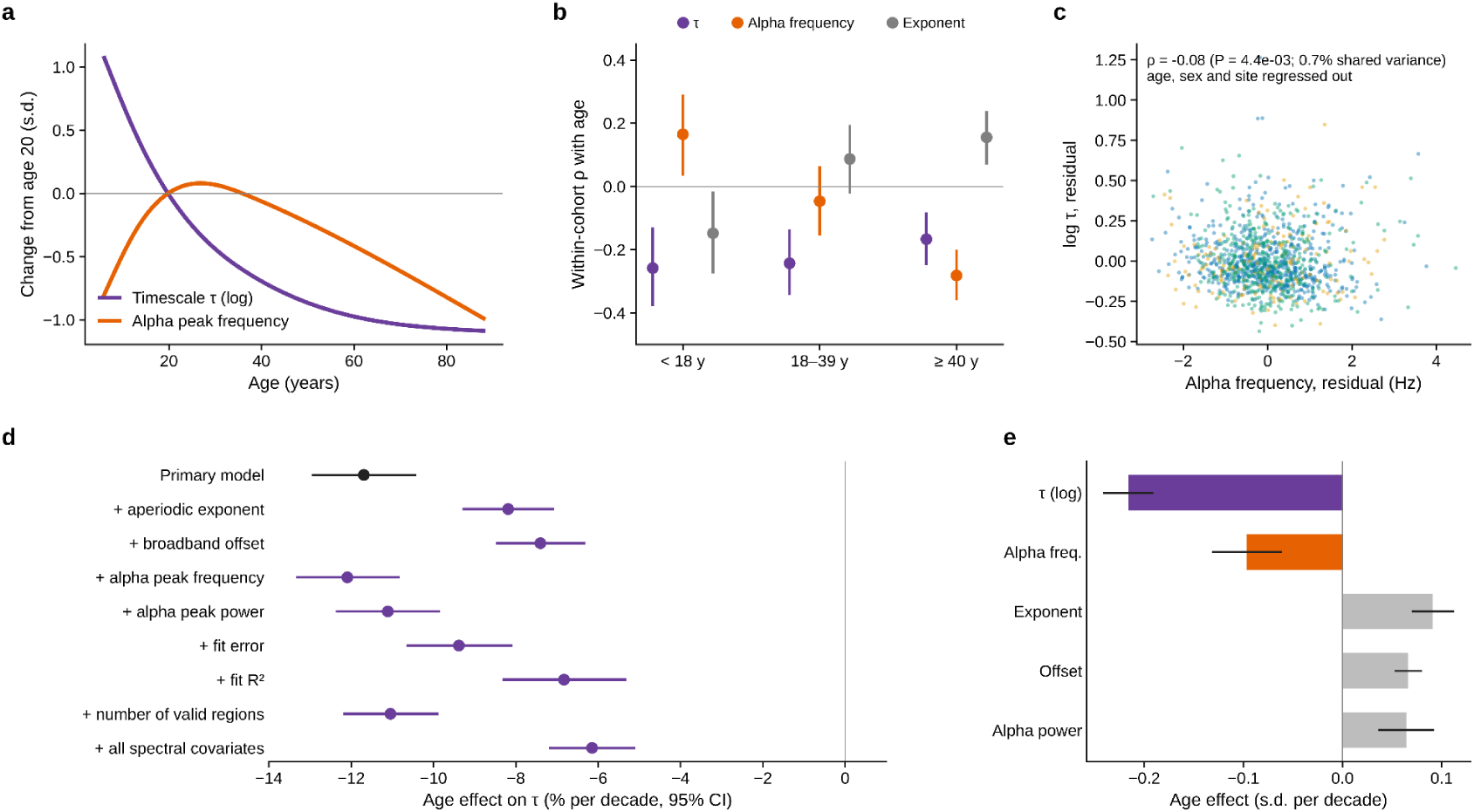
| Intrinsic timescales and alpha oscillations follow distinct lifespan trajectories. **a,** Spline trajectories of log τ (purple) and alpha peak frequency (orange) as standardised change from age 20. τ declines most steeply in development and early adulthood; alpha frequency rises in development, peaks in early adulthood and declines thereafter. **b,** Within-cohort Spearman correlations with age (points; bars, 95% CI) for τ, alpha peak frequency and the aperiodic exponent in three age groups (<18, 18–39 and ≥40 years). **c,** Residual τ against residual alpha peak frequency after regressing out age, sex and cohort (points, participants; colours, cohorts): ρ = −0.08, *P* = 4.4 × 10⁻³, ρ² < 0.01. **d,** Age effect on τ (% per decade; bars, 95% CI) in the primary model and after adjusting for the aperiodic exponent, broadband offset, alpha peak frequency, alpha peak power, spectral fitting error, fit *R*², number of valid regions, or all spectral covariates jointly. The age effect stays negative in every model. **e,** Standardised age effects (s.d. per decade; bars, 95% CI) on log τ, alpha peak frequency, aperiodic exponent, broadband offset and alpha peak power.

The two measures also showed limited correspondence across individuals. After controlling for age, sex and cohort, residual intrinsic timescales and alpha peak frequency were weakly correlated (Spearman ρ = −0.08, ρ² < 0.01; Fig. 4c). Additional adjustment for alpha frequency had little effect on the estimated age association with intrinsic timescales (−11.7% versus −12.1% per decade; Fig. 4d). The standardized age association was also larger for intrinsic timescales than for alpha frequency (−0.22 versus −0.10 standard deviations per decade; Fig. 4e). In comparison, intrinsic timescales were strongly associated with the aperiodic spectral exponent (ρ = −0.82), which is mathematically coupled to the knee parameter within the fitted spectral model. Adjusting for the exponent attenuated the estimated age effect to −8.2% per decade, and jointly adjusting for the measured spectral characteristics further reduced it to −6.2% per decade. Because these covariates partly describe the same spectral structure used to estimate intrinsic timescales, these adjusted associations should not be interpreted as isolating independent physiological mechanisms. Together, the contrasting age trajectories and weak adjusted interindividual association suggest that intrinsic timescales and alpha frequency capture partly dissociable aspects of neural temporal dynamics.

### Cortex-only fields cannot account for the spectra, whereas adding thalamic loops can

Before interpreting the fits, we asked whether thalamic loops are needed at all. A knee fit describes τ but does not explain it, and a cortical field has only fast synaptodendritic and axonal rates and no delayed feedback with which to generate an alpha resonance^15,16^. If the thalamic terms were dispensable, a cortex-only version of the same model should fit the spectra and reproduce their age effects. We tested four nested models (Supplementary Fig. 14; Supplementary Table 9): a cortex-only field (G_ese_ = G_esre_ = G_srs_ = 0; 5 free parameters), the same field with a corticothalamic relay loop and its delay (G_ese_, t_0_; 7), with a reticular pathway added (G_esre_; 8), and the full model with the intrathalamic loop (G_srs_; 9). All were fitted to every participant’s cortex-averaged spectrum with identical objective, bounds and optimizer.

The cortex-only model fitted poorly (median R² = 0.69, interquartile range 0.51–0.80), adding the relay and reticular loops without the intrathalamic loop helped only modestly (R² = 0.76 and 0.75), and the full model reached 0.985 (0.974–0.992). The Bayesian information criterion preferred the full model in 99.9% of participants (median ΔBIC 122 against the cortex-only model). Because adjacent frequency bins are correlated, we also withheld contiguous 9-Hz blocks of the spectrum, refitted each model to the remaining bins and scored the held-out bins (n = 300 participants, five blocks). Median held-out RMSE was 0.141 log_10_ units for the full model (in-sample, 0.049) against 0.328, 0.309 and 0.315 for the cortex-only, relay-loop and reticular-loop models; the full model predicted held-out bins better than the cortex-only model in 98% of participants (paired Wilcoxon P = 3.9 × 10^−50^).

The reduced models also failed qualitatively. A 7–14 Hz spectral peak could be recovered from 15% of cortex-only spectra, compared with 91% of full-model and 92% of MEG spectra, and a valid knee from 15% and 89%, respectively. Where a timescale could be estimated, the cortex-only model did not track individual MEG timescales (ρ = −0.15, 95% CI −0.30 to −0.01), whereas the full model did (ρ = 0.87, 95% CI 0.86 to 0.88). The full model reproduced the age-related shortening of τ in direction and approximate size (model ρ = −0.33, 95% CI −0.38 to −0.27; MEG ρ = −0.42, 95% CI −0.47 to −0.37), whereas the cortex-only model did not (ρ = 0.13, 95% CI −0.02 to 0.27). Within this model family, a delayed corticothalamic loop and an intrathalamic loop are therefore needed to account for these spectra. This does not exclude other cortical architectures or show that the thalamus generates the rhythm; it shows that the thalamic terms are not decorative.

### A corticothalamic model links compression to the intrathalamic loop, and delay explains only part of alpha slowing

To identify candidate circuit-level contributions to the distinct temporal trajectories, we used a nine-parameter corticothalamic neural-field model and estimated its parameters from each participant’s cortex-averaged MEG power spectrum (n = 1,160, including four OMEGA participants without age who contributed to spectral parameter estimation but not to age analyses; median log-spectral R² = 0.985; Fig. 5a; Supplementary Figs. 7 and 8; Supplementary Table 4). These participant-level parameter estimates were used to investigate lifespan associations with intrinsic timescales and alpha frequency. In parallel, we estimated region-specific model parameters from the spectra of individual cortical regions in a cohort- and age-stratified subsample of 280 participants to examine their spatial organization and regional age dependence (Supplementary Figs. 9–11). Intrinsic timescales and alpha peak frequencies derived from the model-predicted spectra closely corresponded to empirical estimates (Spearman ρ = 0.87 and 0.83, respectively; Fig. 5b). The regional parameter estimates revealed spatial variation in corticothalamic dynamics, including associations between intrathalamic feedback gain and cortical intrinsic timescales, as well as age-related changes in selected parameters (Supplementary Figs. 9–11). These regional analyses complemented the participant-level estimates but did not provide independent validation of the inferred physiological mechanisms. Across the three cohorts, the estimated intrathalamic feedback gain Gsrs became more negative with age (within-cohort meta-analytic ρ = −0.48), consistent with stronger effective intrathalamic feedback. The composite thalamic loop parameter Z increased (ρ = 0.42), whereas the synaptodendritic decay rate α decreased (ρ = −0.33) and the estimated corticothalamic propagation delay t₀ increased modestly (ρ = 0.09; Fig. 5c; Supplementary Fig. 11; Supplementary Table 5).

**Fig. 5.**
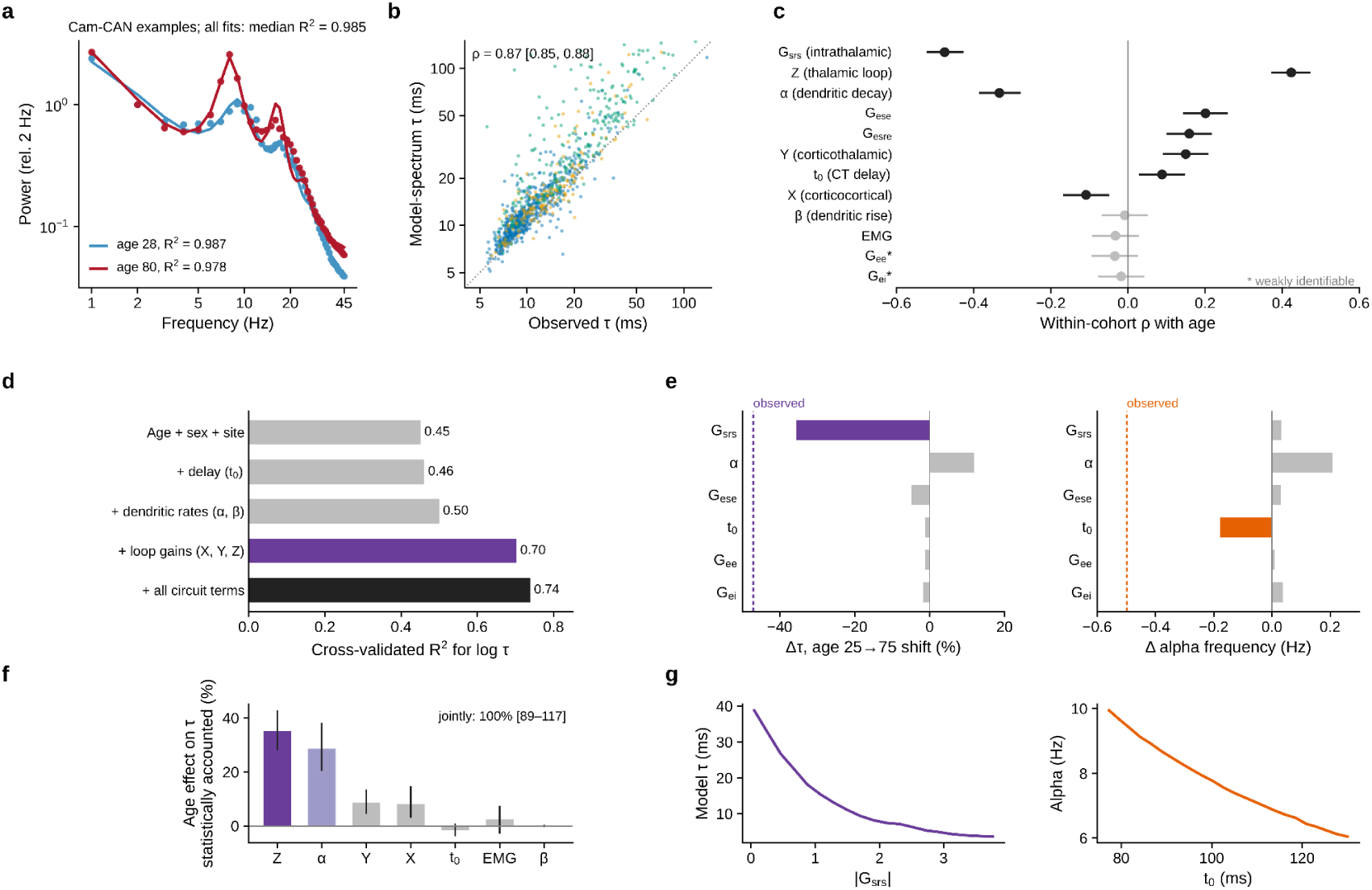
| Individualised corticothalamic models relate timescale compression to intrathalamic dynamics. **a,** Model fits (lines) to the MEG spectra (points) of a 28-year-old and an 80-year-old Cam-CAN participant; the median log-spectral *R*² is 0.985 across all fits. **b,** τ from empirical spectra against τ re-estimated from model spectra (points, participants; dotted line, equality; Spearman ρ = 0.87, 95% CI 0.85–0.88). **c,** Within-cohort Spearman correlations of age with parameters from the cortex-average fits (n = 1,156 with age; points; bars, 95% CI). Grey, not significant after FDR correction. Asterisks mark parameters with limited identifiability (Supplementary Fig. 7). **d,** Cross-validated *R*² for log τ with progressively expanded predictors: age, sex and cohort (0.45); plus *t*_0_ (0.46); plus α and β (0.50); plus loop gains *X*, *Y* and *Z* (0.70); full circuit (0.74). **e,** In silico perturbation: each parameter is shifted from its age-25 to its age-75 value with the others held constant; bars, resulting change in model τ (%, left) and alpha frequency (Hz, right); dashed lines, observed changes. Shifting all parameters together did not reproduce alpha slowing (Supplementary Table 6). **f,** Statistical accounting of the age effect on τ by each fitted parameter (bars, proportion; error bars, uncertainty interval). *Z* and α account for the largest shares and the joint set for about 100% (interval 89–117%). This is statistical accounting, not causal mediation. **g,** Model sweeps of |*G*_srs_| against τ (left) and of *t*_0_ against alpha frequency (right). All model results describe parameter sensitivity within the fitted model and do not establish unique physiological mechanisms.

Fitted circuit parameters accounted for additional interindividual variation in intrinsic timescales beyond age, sex and cohort (cross-validated R² increasing from 0.45 to 0.74; Fig. 5d). A descriptive statistical decomposition attributed 35% of the age association to Z and 29% to α (Fig. 5f). Because the neural timescales and fitted circuit parameters were derived from the same power spectra, these associations quantify shared spectral variation rather than causal mediation or independent predictive validation. We next performed controlled in silico perturbations, shifting individual parameters from their estimated values at age 25 to those at age 75 while holding the remaining parameters constant (Fig. 5e,g; Supplementary Table 6). The age-associated change in G_srs_ shortened model-derived intrinsic timescales by 36%, compared with an empirically estimated reduction of 47%, while producing little change in alpha frequency (+0.03 Hz). Conversely, changing t₀ reduced model alpha frequency by 0.18 Hz, compared with an observed reduction of 0.50 Hz, while leaving intrinsic timescales nearly unchanged (−1%). The remaining parameter changes collectively increased intrinsic timescales by 4%. Shifting all parameters together shortened τ by 37% but raised model alpha frequency by 0.17 Hz, because the age-related decrease in α, which lengthens τ, also raised the alpha peak and offset the effect of delay; this substitution analysis therefore did not reproduce alpha slowing. In the fitted spectra themselves, between 20–29 and 70–89 years the median alpha frequency fell by 0.73 Hz (MEG, 1.31 Hz) and the median τ by 38% (MEG, 45%), so the fitted parameters capture most of the shortening of τ but only about half of alpha slowing (Supplementary Table 6).

### A data-constrained recurrent network contains a slow collective mode that tracks the spectral timescale, unlike single units

The corticothalamic neural-field model allowed us to estimate effective circuit parameters from MEG power spectra and examine their contributions to intrinsic timescales and alpha oscillations. However, spectral modelling alone could not establish how these timescales emerge from interactions among neural populations or how they relate to responses to external inputs. To address these questions, we implemented the fitted corticothalamic dynamics as a time-domain recurrent rate network, preserving the estimated circuit parameters and feedback architecture (Fig. 6a; Supplementary Fig. 12). This complementary implementation allowed us to distinguish individual-unit kinetics from collective network modes and to assess the relationship between spontaneous timescales and stimulus-dependent memory. The network represented the cortical excitatory field, thalamic relay nucleus and thalamic reticular nucleus using interacting chains of leaky units, with time constants, effective coupling strengths and corticothalamic propagation delays derived from the fitted neural-field model. Under the young-adult parameter configuration, individual-unit time constants were no greater than 10.7 ms (about 12.2 ms in the older-adult configuration). The time-domain network reproduced the corresponding analytical model spectra for young-adult and older-adult parameter configurations (Pearson r = 0.997–0.998 for log-transformed spectral power; Fig. 6b), supporting the numerical consistency of the two model implementations.

**Fig. 6.**
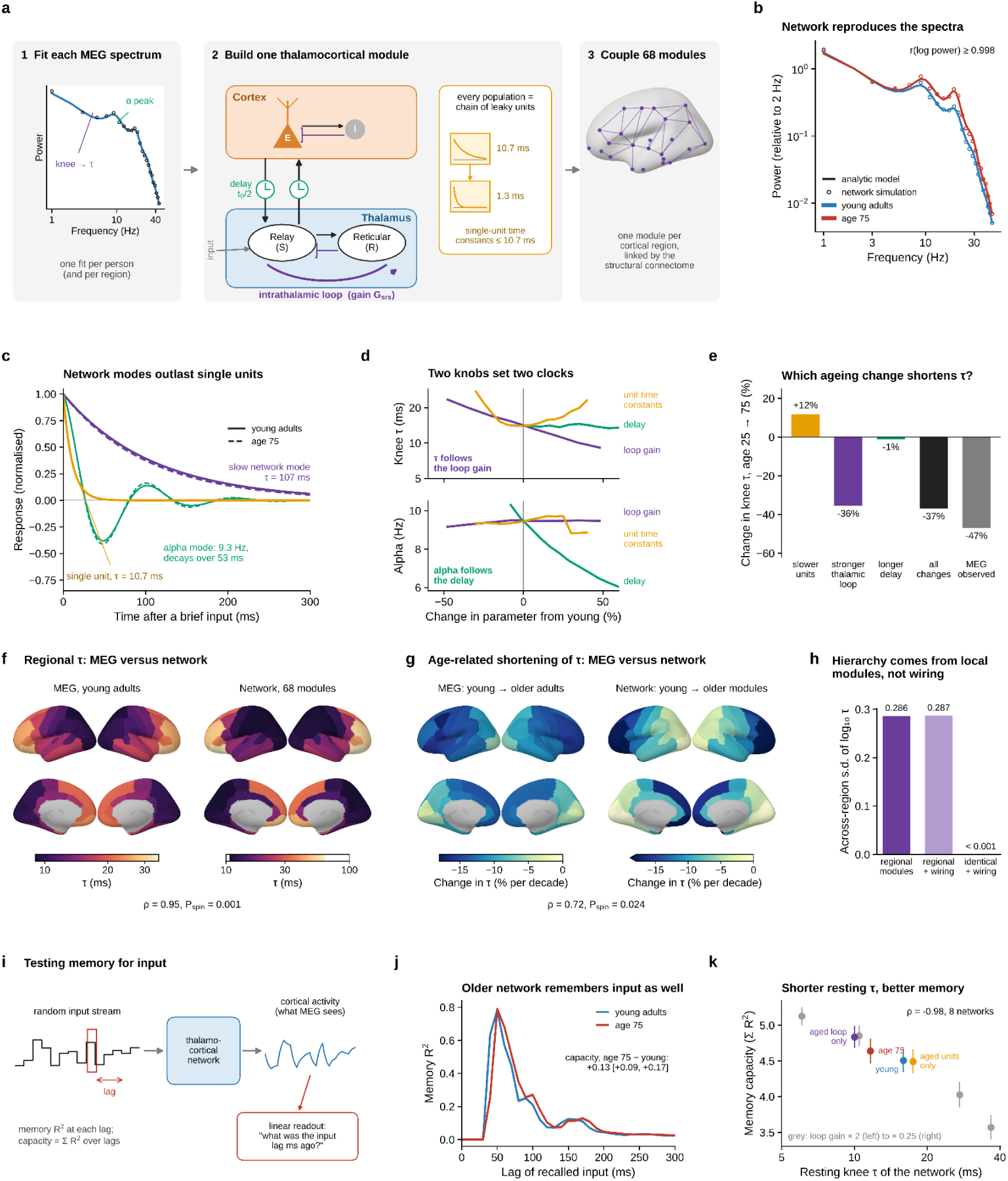
| A data-constrained corticothalamic recurrent network dissociates intrinsic timescales from single-unit dynamics and input memory. **a,** Construction. A neural-field model is fitted to each MEG spectrum (1) and rewritten as a rate network of one thalamocortical module (2): a cortical excitatory field (E; cortical inhibition enters through *G*_ei_), thalamic relay (S) and reticular (R) nuclei, corticothalamic delays of *t*_0_/2 and an intrathalamic loop with gain *G*_srs_; each population is a chain of leaky units (1/α ≈ 10.7 ms, 1/β ≈ 1.3 ms). Sixty-eight modules are coupled by the structural connectome (3). Orange, unit time constants; purple, intrathalamic gain and slow mode; green, delay and alpha mode; blue, young; red, age 75. **b,** Spectra of the time-domain network (circles) and analytical model (lines); Pearson *r* = 0.997–0.998 for log power. **c,** Response to a brief input of the two dominant modes and a single unit: slow mode ≈ 107 ms (102 ms at age 75); alpha mode (9.3 Hz) ≈ 53 ms; both outlast every unit (≤ 10.7 ms young; ≤ 12.2 ms age 75). **d,** Knee τ (top) and alpha frequency (bottom) as |*G*_srs_|, *t*_0_ or unit time constants are varied. **e,** Change in knee τ when each parameter is shifted from its age-25 to its age-75 value, with the MEG-observed change. **f,** Regional τ in young adults (20–39 years): MEG (left) and network (right; ρ = 0.95, *P*_spin_ = 0.001). **g,** Regional age-related shortening (% per decade): MEG (left) and network with older modules (right; ρ = 0.72, *P*_spin_ = 0.024); values below −18% saturate the colour scale. **h,** Across-region s.d. of log_10_ τ for regionally fitted modules, with and without connectome coupling, and for identical coupled modules. **i,** Memory task: random input drives the relay nuclei; a linear readout of cortical activity recalls it at each lag (capacity, summed *R*² over lags). **j,** Memory curves, young and age-75 networks (mean ± s.d.); Δ capacity = +0.13 (95% CI 0.09–0.17). **k,** Capacity against knee τ in eight networks (young, age 75, single-parameter aged networks and *G*_srs_ scaled 0.25–2-fold; ρ = −0.98). Model sensitivities only, not mechanisms.

Eigenmode decomposition of the linearized network revealed a slow, non-oscillatory collective mode with a decay time of approximately 107 ms under the young-adult parameter configuration (102 ms at age 75), substantially longer than the 10.7-ms time constant of its slowest individual units. A separate oscillatory mode occurred at approximately 9.3 Hz, with a decay time of 53 ms. Both modes therefore outlast any single unit when the network responds to a brief input (Fig. 6c; full mode spectrum in Supplementary Fig. 13e). Controlled parameter sweeps showed that increasing effective intrathalamic feedback altered the aperiodic knee timescale in parallel with the slow collective mode (Spearman ρ = 1.0 across the parameter sweep), with comparatively little effect on alpha frequency. The slow mode (about 107 ms) and the spectral knee timescale (about 14 ms) are different quantities, an eigenvalue of the linearised network and an effective timescale estimated from the output spectrum, so we interpret their parallel change as covariation under parameter perturbation and not as identity. Conversely, varying corticothalamic propagation delay shifted alpha frequency from 10.3 to 6.0 Hz while the knee timescale remained between 13.4 and 14.6 ms (Fig. 6d). Age-associated changes in synaptodendritic kinetics lengthened the individual-unit time constants by approximately 14% and, when applied alone, increased the model-derived intrinsic timescale by 12%. In contrast, changing intrathalamic feedback shortened the model-derived timescale by 36% (Fig. 6e; Supplementary Table 6). Within this model, timescale compression can therefore emerge from altered recurrent network dynamics despite a slowing of individual-unit kinetics.

We next extended the model to 68 thalamocortical modules, each corresponding to a Desikan–Killiany cortical region, with region-specific parameters estimated from group-averaged spectra in younger (20–39 years) and older (60–89 years) adults. Modules were coupled using a human structural connectome (Fig. 6f–h; Supplementary Fig. 13a–c). The resulting network closely reproduced the empirical young-adult distribution of regional intrinsic timescales (Spearman ρ = 0.95, P_spin_ = 0.001). Within the tested coupling regime, interregional structural coupling had a relatively small effect on regional timescales, increasing the median by approximately 0.35%. Moreover, structurally coupled modules with identical local parameters did not reproduce the observed cortical hierarchy: the across-region standard deviation of log_10_ τ was 0.286 for regionally fitted modules, 0.287 after connectome coupling and < 0.001 for identical modules coupled through the connectome (Fig. 6h). Replacing younger-adult regional parameters with older-adult estimates shortened intrinsic timescales in 66 of 68 regions, with a median reduction of 8.5% per decade. The spatial pattern of model-derived age effects corresponded to the empirical regional age-effect map (ρ = 0.72, P_spin_ = 0.024), including relatively smaller reductions in visual cortex (−0.4% versus −9.2% per decade). No single parameter change reproduced the complete regional age-effect pattern (Supplementary Fig. 13c; Supplementary Table 7). These simulations suggest that region-specific corticothalamic dynamics are sufficient to reproduce much of the observed spatial organization of intrinsic timescales within this model class, whereas interregional structural coupling makes a comparatively small contribution under the simulated conditions.

### A shorter resting timescale need not mean a shorter memory for input

Finally, we asked whether shorter spontaneous intrinsic timescales necessarily imply reduced capacity to retain or integrate external inputs. Using the 68-module recurrent network as a reservoir, we delivered temporally varying inputs to the thalamic relay populations and trained linear readouts of cortical activity to reconstruct preceding inputs at specified lags (Fig. 6i) or their running averages across integration windows of 10–500 ms (Supplementary Fig. 13d; Supplementary Table 8). Despite exhibiting shorter spontaneous timescales, the older-adult network showed a modest increase in input-memory capacity relative to the young-adult network (Δ = +0.13, 95% CI 0.09–0.17, across eight paired simulation replicates; Fig. 6j). Temporal integration performance over 100–500-ms windows was broadly preserved (Supplementary Fig. 13d). Across eight network configurations with different intrathalamic feedback strengths, shorter spontaneous timescales were associated with greater input-memory capacity (Spearman ρ = −0.98; Fig. 6k). These results indicate that spontaneous neural persistence and stimulus-dependent information retention can vary in different directions within the fitted network. Thus, a shorter resting-state intrinsic timescale does not necessarily imply impaired temporal integration or input memory, although whether this dissociation occurs in the human brain remains to be established through behavioural and physiological validation.

## Discussion

The ageing brain does not simply slow down. Across three cohorts comprising 1,156 participants aged 6–88 years, cortical intrinsic timescales shortened by approximately two-thirds across the lifespan, with negative age associations throughout the cortex and the steepest decline occurring before approximately age 30. In contrast, alpha oscillatory frequency followed a different lifespan trajectory and showed only a weak association with intrinsic timescales across individuals after adjustment for age, sex and cohort. Individualised corticothalamic models associated timescale compression with stronger effective intrathalamic feedback, whereas longer corticothalamic propagation delays partly accounted for alpha slowing within the model. A recurrent network constructed from the fitted parameters further revealed a slow collective mode that covaried with the spectral timescale despite much shorter individual-unit time constants. Together, these findings suggest that the temporal organisation of cortical activity changes across life through partly dissociable dynamical processes rather than a uniform slowing of neural activity.

Previous studies have established that intrinsic timescales increase from sensory to association cortex and are shaped by cortical microarchitecture and recurrent circuit properties^1–3,5,6^. Hierarchical temporal organisation has also been demonstrated at the macroscale using human functional neuroimaging^34^, while earlier lifespan electrophysiological studies reported age-related changes in spontaneous cortical dynamics⁴. Our findings extend this literature in four respects. First, we identify a nonlinear lifespan trajectory in which timescale compression is concentrated in development and early adulthood, with estimated reductions of approximately 30% per decade before a breakpoint around age 30 (95% CI 20–50 years), compared with 6.4% per decade thereafter. Second, although all cortical regions exhibited shorter timescales with increasing age, the magnitude of compression was not systematically organised along the principal functional gradient. The sensory-to-association hierarchy was already present in childhood and remained broadly preserved in later life, with relatively smaller age-related reductions in early visual cortex. Its alignment with the functional gradient was numerically weaker in older adults, although differences between age groups were not significant under spatial permutation. Third, intrinsic timescales and alpha frequency followed distinct age trajectories and showed limited correspondence across individuals. Fourth, computational modelling suggested that local corticothalamic dynamics, rather than interregional structural connectivity alone, were sufficient to reproduce much of the spatial hierarchy and its age-related changes. This distinction is consistent with large-scale modelling studies showing that regional circuit heterogeneity and network connectivity make complementary contributions to cortical dynamics^5,35,36^.

The nested model comparison helped identify which circuit components were required to reproduce the observed spectral features within the corticothalamic neural-field framework. The cortex-only model provided a substantially poorer fit than the full corticothalamic model (median R² = 0.69 versus 0.985), failed to reproduce alpha peaks in most spectra and did not capture the observed age-related shortening of intrinsic timescales. Adding relay and reticular pathways without intrathalamic feedback produced only modest improvements. In contrast, the full model reproduced both the aperiodic knee and the alpha resonance, and performed better in blocked spectral cross-validation. These results suggest that delayed corticothalamic feedback and intrathalamic interactions contribute differently to the organisation of spontaneous neural dynamics, consistent with established roles of thalamocortical circuits in generating and regulating brain rhythms^7–9,15–18^. However, this conclusion is specific to the tested model family and does not establish that the thalamus is the unique source of these dynamics. Both cortical and thalamic generators contribute to the human alpha rhythm^19^, and alternative cortical architectures incorporating recurrent interactions, laminar circuitry or long-range delays could potentially generate similar spectral features^5,6^. The model therefore identifies physiologically plausible circuit mechanisms rather than uniquely determining their anatomical implementation. A testable prediction is that individual differences in thalamic structure, inhibitory signalling or thalamocortical connectivity should be associated with intrinsic timescale variation beyond age alone, particularly given established age-related changes in thalamic anatomy^13,14^.

The functional implications of timescale compression are not straightforward. Intrinsic timescales are commonly interpreted as measures of how long spontaneous neural activity retains information about its preceding state, and hierarchical differences in temporal persistence have been linked to neural processing across multiple temporal scales^1,3,5,34^. Under this interpretation, shorter timescales might imply narrower windows for temporal integration. However, our recurrent-network simulations revealed that shorter spontaneous timescales did not necessarily reduce stimulus-dependent information retention. Instead, increased effective intrathalamic feedback shortened the spectral timescale while preserving or modestly increasing input-memory capacity. This dissociation suggests that spontaneous temporal persistence and the ability to retain externally presented information need not vary in parallel. One possible interpretation is that changes in recurrent feedback reduce the persistence of spontaneous fluctuations while altering the network’s responsiveness to external inputs. Nevertheless, this remains a model-based prediction rather than an experimentally demonstrated developmental mechanism. Direct measurements of temporal integration, working memory and stimulus-evoked neural responses will be needed to establish whether the simulated dissociation occurs in humans. Importantly, the slow collective mode identified in the network and the spectral knee timescale are distinct quantities; their parallel changes under parameter perturbation support a relationship between collective dynamics and spectral timescales but do not establish that they represent the same decay process.

Several observations support the robustness of the lifespan association while also highlighting the challenges of interpreting electrophysiological timescales. The negative age association was present within all three cohorts and remained after accounting for cohort differences, restricting analyses to adults or overlapping age ranges, varying participant inclusion criteria and applying alternative spectral parameterisations. Autocorrelation-based estimates also shortened with age, supporting the conclusion that the effect was not specific to the parametric knee model. Nevertheless, the spectral knee, aperiodic exponent and broadband power are interrelated features of the same neural power spectrum^26^. Previous studies have demonstrated age-related changes in the aperiodic spectral slope, potentially reflecting alterations in the statistical structure of spontaneous neural activity³⁷. Accordingly, the strong relationship between the estimated timescale and aperiodic exponent should not be interpreted as evidence for independent physiological mechanisms. Although adjustment for spectral power, fitting quality and alpha characteristics did not eliminate the age association, changes in source signal quality, cortical atrophy and head-to-sensor distance may still contribute to the observed effects. Moreover, autocorrelation-derived and knee-derived timescales provide related but non-identical estimates of temporal persistence. Thus, the results establish a robust age-related change in the measured temporal characteristics of cortical activity, while the precise physiological processes contributing to this change remain to be determined.

The interpretation of the computational results requires similar caution. The fitted circuit parameters describe effective interactions in a mean-field model and are not direct measurements of thalamic inhibition, synaptic kinetics or axonal conduction. Furthermore, both the fitted parameters and empirical intrinsic timescales were derived from the same MEG power spectra. Their statistical associations and decomposition therefore quantify shared spectral variation rather than independent physiological prediction or causal mediation. Within the model, stronger effective intrathalamic feedback was associated with shorter timescales, whereas longer corticothalamic propagation delays contributed to lower alpha frequency. However, the fitted model reproduced only part of the observed alpha slowing, indicating that additional mechanisms are likely involved. The recurrent-network implementation further demonstrated that collective modes could persist substantially longer than individual-unit time constants and that changing intrathalamic feedback could shorten the spectral timescale despite slower single-unit kinetics. These findings are consistent with theoretical accounts in which neural timescales emerge from recurrent network interactions rather than individual cellular properties alone^3,5,6^. Similarly, the finding that regional circuit heterogeneity reproduced the cortical timescale hierarchy more effectively than interregional structural coupling alone is consistent with models emphasising the importance of spatially heterogeneous local circuit properties³⁵, although anatomical connectivity can shape neural dynamics across multiple temporal scales³⁶. These mechanistic interpretations remain conditional on the model architecture and parameter estimates and require validation using independent physiological measurements or targeted perturbations.

Several limitations should be considered. First, the study was cross-sectional, and age was partially confounded with cohort at the extremes of the lifespan, as all participants younger than 18 years were drawn from PANS. Recording conditions also differed across cohorts, with eyes-open resting-state recordings in OMEGA and eyes-closed recordings in Cam-CAN and PANS. Although age associations were consistently negative within each cohort and robust across cohort-adjusted analyses, longitudinal measurements are needed to establish individual developmental and ageing trajectories. Second, the frequency ranges, spectral estimation procedures and validity criteria may influence the estimated knee timescale, particularly in younger participants whose spectral knees more frequently fell below the lower fitting frequency. Nevertheless, restricting estimates to knees within the fitted frequency range preserved the negative age association. Third, the fitted corticothalamic model contained parameters with limited identifiability, and its superiority was established relative to nested alternatives rather than other neural circuit families. Finally, the recurrent-network analyses provided predictions about collective dynamics and input memory, not direct evidence of human cognitive performance.

In conclusion, cortical intrinsic timescales compress predominantly during development and early adulthood, across the cortical hierarchy and on a lifespan trajectory distinct from alpha-frequency slowing. The convergence of electrophysiological analyses, corticothalamic modelling and recurrent-network simulations suggests that these changes may reflect alterations in collective circuit dynamics rather than a uniform slowing of individual neural processes. More broadly, our findings distinguish the temporal persistence of spontaneous neural activity from oscillatory frequency and stimulus-dependent information retention, providing a framework for understanding how the temporal organisation of the human brain changes across the lifespan.

## Methods

### Participants

We analysed resting-state MEG from three lifespan cohorts collected under local ethics approval with informed consent (and assent from minors). Cam-CAN used a 306-channel Elekta Neuromag VectorView system (1 kHz; about 8.5 min, eyes closed)^20,21^. OMEGA used a 275-channel CTF system (2.4 kHz; ≥5 min, eyes open)^22^. PANS is the lifespan sample of Rempe et al.^4^, recorded with a 306-sensor MEGIN system (1 kHz; 5–8 min, eyes closed); all 388 participants of this sample were analysed, as in the original report. A participant was analysed if age was known and at least one region yielded a valid knee fit (n = 1,156; Supplementary Table 1). Four OMEGA participants without age were excluded from all age-related analyses (n = 1,156) but were retained in the spectral model fits and comparisons (n = 1,160).

### MEG data preprocessing

MEG preprocessing and source-level analyses were performed in Brainstorm (2016 version)^23^, using default settings unless otherwise specified and following established best-practice recommendations^24^. Continuous MEG recordings were first high-pass filtered at 0.3 Hz using a finite impulse response (FIR) filter to remove slow drifts and DC offsets. The raw data were then visually inspected, and channels or time segments contaminated by excessive noise or nonphysiological artifacts were excluded. Power-line interference (50 Hz for Cam-CAN; 60 Hz for OMEGA and PANS) and its harmonics was attenuated using notch filters, followed by band-pass filtering between 0.6 and 280 Hz to retain physiologically relevant neural activity while suppressing residual low- and high-frequency noise. Cardiac and ocular artifacts were identified from simultaneously recorded electrocardiographic (ECG) and electrooculographic (EOG) signals and removed using signal-space projection (SSP). The cleaned continuous recordings were subsequently divided into non-overlapping 30-s epochs. Spectral estimates were computed separately for each epoch and subsequently averaged across epochs to obtain participant-level spectral profiles.

Source reconstruction was performed using each participant’s T1-weighted anatomical MRI. Cortical surfaces and anatomical segmentations were generated with FreeSurfer (v7.4.1)^25^. Individual cortical meshes were resampled to 15,002 vertices and registered to the fsaverage template using spherical surface-based alignment, thereby enabling vertex-wise correspondence across participants. MEG sensor locations were co-registered to individual anatomical space using digitized anatomical landmarks and scalp surface points collected during the MEG recording.

The forward model was constructed using the overlapping-spheres approach. Source-level cortical activity was estimated using a linearly constrained minimum variance (LCMV) beamformer. Data covariance matrices were estimated from the artifact-cleaned epochs, with 5% regularization applied to improve numerical stability. Noise covariance matrices estimated from empty-room recordings were incorporated during source reconstruction to account for sensor noise and reduce spatial biases in the resulting source estimates.

### Neurophysiological spectral calculation

Power spectral density (PSD) was estimated from source-reconstructed magnetoencephalography (MEG) recordings using Welch’s method implemented in Brainstorm^23^. For each recording epoch, PSDs were computed at individual cortical vertices using windows of 2 s (Cam-CAN and OMEGA) or 1 s (PANS) with 50% overlap and subsequently averaged across epochs. Spectral estimates spanned 0 Hz to the Nyquist frequency of each recording, with a frequency resolution of 0.5 Hz (Cam-CAN and OMEGA) or 1 Hz (PANS). Vertex-level spectra were then aggregated into 68 cortical regions defined by the Desikan–Killiany atlas on the FreeSurfer FSAverage template. Within each region, principal component analysis (PCA) was applied to the vertex-level spectral profiles, and the first principal component was retained as the representative regional spectrum, with its sign oriented to ensure a positive correlation with the corresponding parcel-averaged spectrum.

To harmonize spectral representations across cohorts, frequency axes were reconstructed from the number of spectral bins and the Nyquist frequency of each cohort-specific Welch estimate, yielding frequency increments of 0.5 Hz for Cam-CAN and OMEGA and 1 Hz for PANS. Frequency assignments were validated against identifiable spectral landmarks, including alpha-band peaks, line-noise notches (50 Hz for Cam-CAN and 60 Hz for OMEGA and PANS), and head-position-indicator peaks. Spectra sampled at 0.5-Hz intervals were smoothed using a three-point kernel [0.25, 0.5, 0.25], after which all regional spectra were resampled onto a common frequency grid spanning 1–100 Hz in 1-Hz increments. Cortical regions were ordered according to Brainstorm’s alphabetical convention, with left- and right-hemisphere parcels interleaved. The number of bins of each cohort matched its sampling rate and window length (1,001 bins to 500 Hz, 2,401 bins to 1,200 Hz and 501 bins to 500 Hz), which is consistent with estimation at the native sampling rate without further resampling or zero-padding. This ordering was supported by the expected decline in interregional spectral similarity with increasing anatomical distance (Spearman’s ρ = 0.57, 0.66 and 0.76 across the three cohorts; Supplementary Fig. 1c). The resulting harmonized regional spectra served as the input for all subsequent analyses.

### Spectral parameterisation and timescale

Each region × participant spectrum was fitted with FOOOF 1.1^26^ in knee mode over 2–45 Hz (peak width 1–12 Hz, ≤6 peaks, minimum height 0.05): log₁₀P(f) = b − log₁₀(k + f^χ). We defined *f*_k_ = *k*^1/χ^ and τ = 1/(2π*f*_k_). Alpha frequency and power were taken from the largest peak in 7–14 Hz. A fit was valid if k > 0, 0 < τ < 500 ms and R² ≥ 0.9; because τ < 500 ms corresponds to *f*_k_ > 0.32 Hz, this criterion does not require the knee to lie inside the fitted band. Of 75,670 valid regional fits, 95.0% had *f*_k_ within 2–45 Hz, 5.0% below 2 Hz and fewer than 0.1% above 45 Hz (by cohort, Cam-CAN 98.3%, OMEGA 96.0%, PANS 86.2%; by age, 82.6% below 18 years, 96.5% at 18–59 years and 98.0% at 60 years or older; Supplementary Results). Participant values are medians over valid regions; statistics used log₁₀ τ. Autocorrelation-based timescales, which do not assume a parametric knee model, were computed from the same spectra as ACF(t) = Σ_f_ P(f) cos(2πft) over 1–45 Hz (sampled every 0.1 ms); τ_ACF_ is the first 1/e crossing and τ_int_ the area under the ACF up to its first zero crossing.

### Corticothalamic neural-field model fitting

We fitted the corticothalamic neural-field model of Robinson and colleagues^15^, re-implemented in Python following the full model of the BrainTrak framework^17^, to the MEG power spectra. The model describes cortical excitatory and inhibitory populations interacting with the thalamic relay and reticular nuclei and has nine free parameters: the cortical excitatory and inhibitory gains G_ee_ and G_ei_, the corticothalamic loop gains G_ese_ and G_esre_, the intrathalamic loop gain G_srs_, the synaptodendritic rate constants α and β, the corticothalamic propagation delay t_0_ and the amplitude of an electromyographic component (equations in Supplementary Methods).

Three sets of fits were obtained. (i) Participant-level fits: for every recording (n = 1,160; 1,156 with age and sex), the target was the cortex-average spectrum (the mean of log_10_ power over the 68 regions, 1–45 Hz), normalized to unit area so that only spectral shape constrained the parameters; these fits provide all participant-level parameter estimates. (ii) Regional fits: the 68 regional spectra of a cohort- and age-stratified subsample of 280 participants (Cam-CAN 100, OMEGA 80, PANS 100) were fitted separately to examine the spatial organization of the parameters. (iii) Group-regional fits: the cohort-balanced mean log spectrum of each region in younger (20–39 years) and older (60–89 years) adults was fitted to parameterize the 68-module recurrent network.

The model was fitted by minimizing a frequency-weighted squared relative error between predicted and empirical power spectra, with inverse-frequency weighting to balance contributions across frequencies. Parameters were constrained to physiologically plausible ranges and required to satisfy linear-stability criteria. Each fit was initialized from the three best candidates in a library of 6,000 valid and stable parameter sets and refined using bounded Nelder–Mead optimization. The solution with the lowest fitting error was retained.

Fitted physiological parameters were transformed into the canonical X, Y and Z coordinates of the BrainTrak neural-field framework to summarize effective corticocortical, corticothalamic and intrathalamic feedback. Model fit quality was evaluated using the root-mean-square error (RMSE) and coefficient of determination (R²) between empirical and predicted log-transformed power spectra. Unsuccessful fits were excluded from parameter-based analyses. Full model equations, parameter transformations, fitting procedures, parameter bounds, identifiability assessments and test–retest reproducibility are detailed in the Supplementary Methods, Supplementary Figs. 7 and 8 and Supplementary Table 4.

### Assessment of model-fitting quality

To determine whether model fit varied systematically across individuals, the log-spectral R² of the participant-level fits was related to age within each cohort and summarized by cohort (Supplementary Results).

### Nested model comparison

To test whether thalamic terms are needed, we fitted four nested versions of the model, each obtained by fixing subsets of the nine parameters to zero. The cortex-only model fixed G_ese_, G_esre_ and G_srs_ to zero and left G_ee_, G_ei_, α, β and EMG_a_ free (5 free parameters). The relay-loop model additionally freed G_ese_ and t_0_ (7), the reticular-pathway model freed G_esre_ (8), and the full model freed G_srs_ (9). All four used the same equations, parameter bounds, stability and validity rules, objective (χ² with w(f) = 1/f), 6,000-set stable starting library, three best starting points and bounded Nelder–Mead refinement as the main fits (1–45 Hz, 45 bins). The model spectrum was scaled to the target by total power over the fitted bins. Every participant (n = 1,160: the 1,156 participants of the primary analysis and four OMEGA participants without age, who contribute to model fitting but not to age analyses) was fitted with every model.

Fit quality was summarized as R² and RMSE of log_10_ power, and the Bayesian information criterion BIC = n ln(RSS/n) + k ln n, with n = 45 bins, RSS the residual sum of squares of log_10_ power and k the number of free parameters. Because adjacent bins are correlated, BIC was used descriptively. Out-of-sample performance was assessed by blocked cross-validation: the 45 bins were divided into five contiguous 9-Hz blocks, each model was refitted to the remaining 36 bins, and the mean squared error of log_10_ power was computed on the held-out block. This was done for 300 participants (100 per cohort, evenly spaced across the age-ranked participants of each cohort) and the five block errors were averaged for each participant. Models were compared with paired Wilcoxon signed-rank tests across participants. Timescales and alpha peak frequencies of model spectra were re-estimated with the same spectral-parameterization settings and validity criteria (knee > 0, τ < 500 ms, R² ≥ 0.9) used for the MEG spectra, and were related to the empirical values and to age by within-cohort Spearman correlation pooled by inverse-variance weighting on the Fisher z scale. Fits at a parameter bound were identified with a tolerance of 1% of the parameter range.

### Thalamocortical recurrent network

To examine whether the spectral timescale could emerge from collective network dynamics rather than single-unit kinetics, we converted the fitted corticothalamic neural-field model into a time-domain recurrent rate network. Each thalamocortical module comprised interacting cortical excitatory, thalamic relay and reticular populations implemented as chains of leaky units, with time constants, effective coupling strengths and corticothalamic delays derived directly from the fitted neural-field parameters. We first analysed a single module using eigenmode decomposition to quantify collective decay times and oscillatory frequencies. We then constructed a 68-module network using region-specific fitted parameters and coupled the modules through the group-average structural connectome. Finally, we used this network as a reservoir to test whether spontaneous intrinsic timescales predicted the capacity to retain and integrate temporally varying inputs. Full network equations, numerical implementation and simulation procedures are provided in the Supplementary Methods.

### Statistical accounting and in silico perturbation

Cross-validated R² was computed for log_10_ τ of the cortex-average MEG spectrum (n = 1156 participants with age and a valid τ) using 10-fold cross-validation repeated 20 times. The baseline model contained age in decades (linear and quadratic terms), sex and cohort; circuit terms (X, Y, Z, α, β, t_0_, EMG_a_), each standardised within cohort, were added as indicated in Fig. 5d. The statistical decomposition of the age–τ association used a product-of-coefficients approach. For each fitted quantity Q (standardised within cohort), the indirect effect was a × b, where a is the coefficient of age (in decades) in a regression of Q on age, sex and cohort, and b is the coefficient of Q in a regression of log_10_ τ on Q, age, sex and cohort. The proportion accounted for was a × b divided by the total age effect on log_10_ τ (−0.055 per decade), and the joint proportion was one minus the ratio of the direct age effect to the total effect when all seven circuit terms were included together. Intervals are 95% percentile intervals from 1,000 bootstrap resamples of participants. Because τ and the circuit parameters were derived from the same spectra, this decomposition describes shared spectral variation and does not support causal mediation. For the in silico perturbations, the baseline was the median of each of the nine fitted parameters in participants aged 20–29 years. The value at age 75 of each parameter was the baseline plus the slope of its linear regression on age (with sex and cohort as covariates) multiplied by 50 years, clipped to the parameter bounds; parameters already at a bound in most fits (β and G_esre_) therefore changed little. The model spectrum was recomputed for each configuration, and τ and alpha frequency were extracted with the same spectral-parameterization procedure as for the MEG data. Because each parameter is shifted independently of the others, this analysis ignores correlations among parameters and non-linear interactions; we therefore also compared the age change in τ and alpha frequency of the fitted model spectra themselves with that of the MEG spectra (group medians at 20–29 and 70–89 years).

### Functional gradient and structural connectome

The principal functional gradient was the first diffusion-map component^27^ of the group-average resting-state functional connectivity matrix of the Human Connectome Project^28^ in the Desikan–Killiany parcellation, obtained from the ENIGMA Toolbox^29^; it orders regions from unimodal (visual) to transmodal cortex. The structural connectome used to couple network modules was the corresponding group-average Human Connectome Project structural connectivity matrix (log(1 + streamline density), scaled to unit spectral radius).

### Statistics

Age associations were estimated within cohort (Spearman ρ) and combined by fixed-effect Fisher-z meta-analysis (weights n − 3) with I². Pooled models were ordinary least squares with cohort and sex as fixed effects and HC3 standard errors, age in decades centred at 40 years; effects on log τ are reported as % change per decade. Linear, log-age, quadratic and natural cubic spline (4 d.f.) models were compared by AIC and 10-fold cross-validated R². The breakpoint of a two-segment model was found by profile AIC over 20–65 years with a 300-sample bootstrap. Regional effects were corrected for multiple comparisons over 68 regions with the Benjamini–Hochberg false discovery rate^30^. All statistical tests were two-sided unless stated otherwise. No statistical method was used to predetermine sample size; all participants of the three datasets meeting the inclusion criteria were analysed. Data collection and analysis were not performed blind to participant age, which was the variable of interest.

### Spatial analyses and life-stage maps

Life-stage maps (<18, 18–59, ≥60 years and all ages) were computed as the mean over cohorts (with ≥10 participants in that stage) of the within-cohort regional median of log τ. Maps were rendered on the fsaverage5 inflated surface with ENIGMA toolbox labels ^29^. The hierarchy axis was the first diffusion-map component of the HCP group functional connectivity matrix ^27^. Map correlations were tested against 2,000 spatial rotations of parcel centroids (spin test^31^; 1,000 rotations for the maps generated by the recurrent network).

### Sensitivity analyses

The global age effect was re-estimated across 40 analytic specifications (the primary analysis and 39 variants; Supplementary Table 2): participant inclusion (censored median treating knee-less regions as τ above range; ≥10, ≥34, ≥50 and 68 valid regions), τ thresholds and knee-in-range rules, fit-quality rules, covariates (exponent, offset, alpha frequency and power, fit error and R², number of valid regions, all jointly), cohort handling (single cohort, leave-one-out, ComBat with age and sex protected^32,33^), robust Huber regression, age ranges, sex strata and three FOOOF refits (1–45 Hz, 3–40 Hz, ≤3 peaks of width 2–8 Hz).

## Data availability

MEG and structural MRI data of the Cambridge Centre for Ageing and Neuroscience (Cam-CAN) cohort are available at https://cam-can.mrc-cbu.cam.ac.uk/dataset/. The Open MEG Archive (OMEGA) is available at https://www.mcgill.ca/bic/neuroinformatics/omega. The PANS lifespan dataset^4^ is available from https://cdn.boystown.org/media/Rempe_Ott_PNAS_2023_Data.zip. Group-average Human Connectome Project connectivity matrices were obtained from the ENIGMA Toolbox (https://enigma-toolbox.readthedocs.io/en/latest/pages.html); BrainSpace is available at https://brainspace.readthedocs.io/en/latest/. Source data for all figures are provided with this paper.

## Code availability

All code used in this study is available at https://github.com/Laoma29/MEG_Timescale_networ.

## Acknowledgements

This work was supported by the National Natural Science Foundation of China (grants number 82302175 and 62336002) and the National Science and Technology Innovation 2030 Program (grant number 2021ZD0200506 and 2021ZD0200500). We thank the National Center for Protein Sciences at Peking University in Beijing, China for assistance on data analysis. This work was partially supported by Beijing Institute of Technology Kunpeng&Ascend Center of Cultivation.

## Author contributions

Xiaobo Liu conceived and designed the study, developed the methodology, performed all data processing and analyses, interpreted the results, prepared the figures, and wrote the manuscript. Brendan Harris, Runhao Lu, Sanwang Wang, Qiuxuan Yu, Mingyue Zhang, Ruifang Cui, Xinyu Liang, and Tengteng Fan provided scientific feedback and critically reviewed and revised the manuscript. All authors reviewed and approved the final manuscript.

## Competing interests

The authors declare no competing interests.

## Supplementary Information

### Cortical intrinsic timescales compress early in life, on a schedule separate from alpha slowing

#### Datasets and data acquisition

##### Cambridge Centre for Ageing and Neuroscience Dataset

We analyzed resting-state magnetoencephalography (MEG) data from 606 healthy adults (299 females; mean age = 54.2 ± 18.3 years) from the Cambridge Centre for Ageing and Neuroscience (Cam-CAN) cohort^20^. Cam-CAN is a population-based, cross-sectional lifespan study designed to characterize neurocognitive changes across healthy adulthood. All participants included in the present study had usable resting-state MEG recordings and corresponding structural MRI data.

##### MEG Data Acquisition

MEG data were acquired at the MRC Cognition and Brain Sciences Unit (MRC-CBSU, Cambridge, UK) using a 306-channel VectorView MEG system (Elekta Neuromag, Helsinki, Finland), comprising 102 magnetometers and 204 orthogonal planar gradiometers. Recordings were obtained with participants seated inside a magnetically shielded room. Continuous MEG signals were sampled at 1,000 Hz with an acquisition bandwidth of 0.03–330 Hz.

Head position within the MEG helmet was continuously monitored using four Head Position Indicator (HPI) coils, enabling subsequent correction for head motion. Horizontal and vertical electrooculographic signals were recorded using bipolar electrodes to monitor ocular artifacts, and electrocardiographic signals were recorded simultaneously to characterize cardiac artifacts. During the resting-state session, participants were instructed to remain still, relaxed, awake, and with their eyes closed. Approximately 8 min and 40 s of resting-state MEG data were acquired for each participant.

##### MRI Data Acquisition

High-resolution structural MRI data were acquired during a separate session using a 3 T Siemens TIM Trio scanner equipped with a 32-channel head coil. T1-weighted anatomical images were acquired using a three-dimensional magnetization-prepared rapid gradient-echo (MPRAGE) sequence at 1-mm isotropic resolution. Individual anatomical images were used for cortical surface reconstruction, anatomical parcellation, and MEG-to-MRI coregistration.

##### Open MEG Archives

We independently replicated the principal findings using resting-state MEG data from the Open MEG Archives (OMEGA)^22^. OMEGA is an open-access repository of multimodal neuroimaging data acquired at the Montreal Neurological Institute. The present analysis included 162 unrelated participants (80 females; mean age = 31.6 ± 14.6 years) with usable resting-state MEG recordings.

MEG data were acquired using a 275-channel whole-head CTF system (CTF Systems Inc., Port Coquitlam, British Columbia, Canada). Signals were sampled at 2,400 Hz with an anti-aliasing low-pass filter at 600 Hz, and built-in third-order spatial gradient compensation was applied to attenuate environmental magnetic noise. Approximately 5 min of continuous eyes-open resting-state MEG data were collected while participants remained still and fixated on a centrally presented visual target.

Individual head geometry and fiducial landmarks were digitized using a Polhemus system to facilitate MEG-to-MRI coregistration. Structural T1-weighted MRI data were available for anatomical reconstruction and source localization. High-resolution T1-weighted images were acquired using a three-dimensional MPRAGE sequence with approximately 1-mm isotropic spatial resolution.

A subset of the OMEGA cohort underwent repeated MEG recordings on separate days. Where test–retest analyses were performed, only participants with at least two independent recording sessions were included, and two sessions from each participant were used to quantify between-session reliability.

##### Lifespan MEG Dataset (PNAS Dataset)

To further assess the generalizability of the findings across a substantially broader developmental and ageing range, we analyzed an independent publicly available lifespan MEG dataset originally reported by Rempe et al.^4^. The dataset comprised 388 neurologically healthy participants (183 females) between 6 and 85 years of age (mean age = 25.3 ± 20.9 years). Participants were pooled across five independent studies conducted at Boys Town National Research Hospital, with all MEG recordings acquired using the same MEG instrument and harmonized acquisition procedures.

Adult participants were neurologically and cognitively healthy control participants recruited in studies of neurological or psychiatric conditions, whereas pediatric participants were recruited through studies of typical neurodevelopment. Exclusion criteria in the original studies included neurological or psychiatric disorders, medical conditions affecting the central nervous system, history of significant head trauma, current substance misuse, inability to complete the resting-state recording, and standard contraindications to MEG acquisition. All original studies were approved by the local Institutional Review Board, with written informed consent obtained from adult participants or parents/legal guardians and assent obtained from pediatric participants when appropriate.

##### MEG Data Acquisition

Resting-state MEG recordings were acquired in a magnetically shielded room using a 306-channel MEGIN system (MEGIN, Helsinki, Finland), comprising 102 magnetometers and 204 planar gradiometers. Neuromagnetic signals were continuously sampled at 1,000 Hz with an acquisition bandwidth of 0.1–330 Hz. Participants were seated comfortably inside the MEG system and instructed to remain relaxed with their eyes closed. Resting-state recordings lasted approximately 5–8 min, during which participants were continuously monitored using an audio–video system.

Four head-position indicator coils were attached to the participant’s head and localized together with anatomical fiducials and scalp surface points using a three-dimensional digitizer. These measurements enabled alignment of MEG sensor coordinates with individual anatomical space and subsequent source-level analyses.

##### MRI Data Acquisition and MEG–MRI Coregistration

Structural MRI data were available for anatomical reconstruction and MEG–MRI coregistration. Individual MEG recordings were aligned to structural anatomy using digitized fiducial landmarks and scalp surface points. For participants with individual structural MRI data, subject-specific anatomy was used for source reconstruction. The publicly released dataset includes de-identified MEG recordings, structural MRI data, and demographic information, including participant age and sex.

Together, these three independent datasets provided complementary coverage of the adult lifespan and broader developmental-to-ageing trajectory, while also spanning different MEG systems, acquisition environments, and resting-state protocols. This multi-cohort design enabled us to evaluate the reproducibility and generalizability of the observed cortical dynamics across independent populations and acquisition settings.

##### Supplementary Methods: corticothalamic neural-field modelling

###### Model

Power spectra were modelled with the linearised corticothalamic neural-field model of Robinson and colleagues^15^, as implemented in BrainTrak^17^. The model comprises cortical excitatory and inhibitory populations, thalamic relay (S) and reticular (R) nuclei, and a cortex–thalamus round-trip delay *t*_0_. Synaptodendritic dynamics are described by the low-pass filter

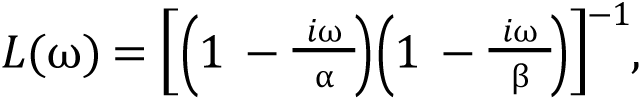

where α and β are the decay and rise rates. The fitted parameters are the cortical gains *G*_ee_ and *G*_ei_, the loop gains *G*_ese_ (cortex–relay), *G*_esre_ (cortex–reticular–relay) and *G*_srs_ (intrathalamic), the rates α and β, the delay *t*_0_, and the amplitude EMG_a_ of an electromyographic component. The cortical dispersion relation is

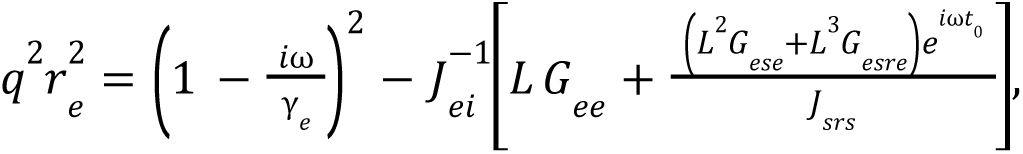

with *J*_ei_ = 1 − *L G*_ei_ and *J*_srs_ = 1 − *L*² *G*_srs_. The power spectrum is

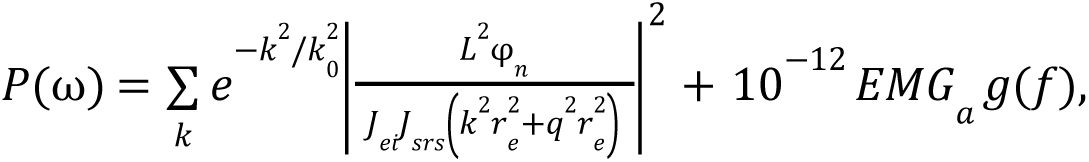

where *g* = *x*²/(1 + *x*²)², *x* = *f*/40 Hz, and the sum runs over spatial modes *k* = 2π*m*/*L*_x_ (|*m*| ≤ 4; *L*_x_ = 0.5 m; *k*_0_ = 10 m⁻¹; γ_e_ = 116 s⁻¹; *r*_e_ = 0.086 m). The spectrum was normalised to the area of the target spectrum, so that only spectral shape constrains the parameters. We also report the composite loop gains *X* = *G*_ee_/(1 − *G*_ei_), *Y* = (*G*_ese_ + *G*_esre_)/[(1 − *G*_srs_)(1 − *G*_ei_)] and *Z* = −*G*_srs_αβ/(α + β)².

###### Spectral target and objective

For each recording (n = 1,160; Cam-CAN 606, OMEGA 166, PANS 388; 1,156 with age and sex), the target was the cortex-average spectrum, obtained by averaging log_10_ power over the 68 Desikan–Killiany regions and exponentiating, restricted to 1–45 Hz (45 frequency bins). The objective was χ² = Σf w(f)[(Pmodel − Pdata)/Pdata]², with w(f) = 1/f, so that each frequency decade contributes approximately equally. Region-level parameters were obtained from separate fits to the 68 regional spectra of a stratified subsample of 280 participants (spatial analyses; Supplementary Figs. 9, 10 and 11) and from two group-level fits (younger and older groups, mean ages 28.4 and 69.7 y; 68 regions each) that parameterized the recurrent network.

###### Constraints

Parameters were bounded as follows: G_ee_ [0, 20], G_ei_ [−40, 0], G_ese_ [0, 40], G_esre_ [−40, 0], G_srs_ [−5, 0], α [10, 100] s⁻¹, β [100, 800] s⁻¹, t_0_ [75, 140] ms, EMG_a_ [0, 1]. Parameter sets were rejected if G_ee_/G_ei_ ≤ −0.5, G_ee_ + G_ei_ > 1, or β/α > 20, and if the k = 0 dispersion relation failed a Nyquist-type linear-stability criterion evaluated at 1,001 frequencies between 0 and 100 Hz.

###### Optimisation

A library of 6,000 valid and stable parameter sets was drawn uniformly within the bounds (EMG_a_ = u³, u ∼ U(0, 1)). The three library members with the lowest χ² were used as starting points for bounded Nelder–Mead optimisation in the unit-cube parameterisation (adaptive simplex, up to 600 function evaluations per start, xatol = 10⁻⁴, fatol = 10⁻⁶; invalid or unstable points were assigned χ² = 10⁶). The solution with the lowest χ² was retained. Goodness of fit was the coefficient of determination of the log_10_ spectrum (log-fit R²).

###### Audit of the fitting procedure

To verify reproducibility, 200 randomly selected recordings were re-fitted with the released code. Median χ² differed from the stored value by 0.5%. The random library alone gave a median χ² of 0.209, which fell to 0.048 after refinement from one start and 0.043 from three starts. Re-fitting 100 recordings with an independently drawn library yielded virtually identical χ² (median ratio 1.001).

###### Identifiability

We used three complementary analyses. (i) Local sensitivity: the Jacobian of the normalised log_10_ spectrum with respect to each parameter (expressed in the unit-cube coordinate) at the young-adult median parameter set, weighted by the objective, and its singular-value decomposition. (ii) Repeatability: Spearman correlation between parameters estimated with two independent libraries (n = 100) and between independent re-fits (n = 60). (iii) Parameter recovery: spectra were generated from 150 fitted parameter sets, perturbed with multiplicative log-normal noise matching the empirical residual (RMS 0.059 log_10_ units), and re-fitted with the complete pipeline. Recovery was quantified as the Spearman correlation between generating and recovered parameters.

###### Age effects

For each parameter, Spearman correlation with age was computed within each cohort and pooled across cohorts by inverse-variance weighting on the Fisher z scale (weights n − 3), with heterogeneity quantified by I². P values were corrected across parameters by the Benjamini–Hochberg procedure.

##### Supplementary Methods: nested neural-field model comparison

The four nested models (M0–M3) were generated from the full model by fixing parameters to zero while leaving all equations unchanged (Methods). In M0, G_ese_, G_esre_ and G_srs_ were zero, so the thalamic relay and reticular populations did not feed back on cortex and the dispersion relation reduced to the cortical field with the local gains G_ee_ and G_ei_ and synaptodendritic filtering by α and β. In M1, the relay loop (G_ese_) and its delay (t_0_) were released; in M2 the reticular pathway (G_esre_) was released as well; and in M3 the intrathalamic loop (G_srs_) was also released. Because the sets of free parameters differed, each model had its own stable starting library of 6,000 parameter sets sampled uniformly within the bounds of its free parameters, with the same stability and validity rules as in the main fits. For blocked cross-validation, the library screening and refinement used only the training bins, and the model spectrum was scaled to the target by total power over the training bins, so that no information from the held-out bins entered the fit. Information criteria were computed from the residual sum of squares of log_10_ power over 45 bins and are shown for description only.

##### Supplementary Methods: thalamocortical rate network

###### Construction

To examine the timescales implied by the fitted circuit in the time domain, we implemented the k = 0 component of the model as a discrete-time rate network with no free connectivity parameters. Each population (cortical E, relay S, reticular R) comprises two leaky units in series with time constants 1/α and 1/β; the cortical field ψ is obtained through two further units with time constant 1/γ_e_. Units were updated by exponential Euler integration, x(t + Δt) = a x(t) + (1 − a) u(t), a = exp(−Δt/τ). Cortico-thalamic and thalamo-cortical projections carried a delay of t_0_/2 implemented as a delay line of D = t_0_/(2Δt) steps. The products of gains in the field model were factorised into edge weights, G_es_ = G_ese_, G_se_ = 1, G_sr_ = −1, G_rs_ = −G_srs_ and G_re_ = −G_esre_/G_ese_, so that, for φ equal to the identity, the transfer function from noise injected into the relay nucleus to ψ equals the k = 0 term of the analytic spectrum. For task simulations, φ = tanh, which has unit slope at the operating point and therefore leaves the linearisation unchanged. Parameter values for the young-adult network (median of 20–29 y) were G_ee_ = 13.58, G_ei_ = −23.32, G_es_ = 7.48, G_rs_ = 1.20, G_re_ = 0.005, 1/α = 10.7 ms, 1/β = 1.25 ms, 1/γ_e_ = 8.62 ms and t_0_/2 = 40.7 ms; the 75-y network was obtained by adding the regression slope on age multiplied by 50 y to each parameter (G_es_ = 9.98, G_rs_ = 1.80, 1/α = 12.2 ms). Because G_esre_ was at its bound in most fits, it was constrained to the interior of the parameter range (−0.04) before conversion to G_re_.

###### Network of modules

For region-level simulations, 68 modules, each with its own region-specific parameters, were coupled through the structural connectome, ue,j += c Σk Wjk ψk. The coupling c was set to half the critical value (c = 2.58; c_crit_ = 5.15, at which the spectral radius of the linearised network reaches 1). The contribution of local parameters versus structural coupling to the timescale hierarchy (Fig. 6h) was quantified as the across-region standard deviation of log_10_ τ for regionally fitted modules simulated in isolation and with connectome coupling, and for identical (young-adult median) modules coupled through the connectome. Spatial correspondence of network-generated maps was assessed with 1,000 spin rotations.

###### Timescale analysis and validation

Network timescales were obtained from the eigenvalues λi of the linearised state-transition matrix, τi = −Δt/ln|λi| and fi = |arg λi|/(2πΔt). Time-domain simulations (4 replicates of 60 s, Δt = 0.25 ms, independent unit-variance noise to the relay nucleus) were compared with the analytic spectra (Pearson correlation of log power) and with timescales estimated from the autocorrelation function. Numerical convergence was evaluated at Δt = 1, 0.5, 0.25 and 0.125 ms. To dissociate unit-level and network-level contributions to the spectral timescale, we varied, one at a time, the unit time constants (×0.7–1.4), G_srs_ (−2.6 to −0.3) and t_0_ (75–130 ms). For display (Fig. 6c), the impulse responses of the two dominant modes were drawn as exp(−t/τ) for the slow real mode and exp(−t/τ)cos(2πft) for the alpha mode, using the eigenvalue-derived decay times and frequencies, and compared with the single-unit decay exp(−tα).

###### Information-processing simulations

A random stream u (uniform on [−1, 1], 10 ms bins) was delivered to the relay nuclei of all 68 modules with module-specific weights (N(1, 0.3), truncated at 0.2), together with independent intrinsic noise (σ = 0.5), for 120 s (Δt = 0.5 ms; first 2 s discarded). A ridge-regression readout (regularisation chosen by cross-validation over 10⁻³–10³) was trained on the z-scored cortical activity ψ of the 68 modules in the first 70% of the data and evaluated in the last 30%. Two tasks were assessed with the cross-validated R²: recall of u at lags of 0–400 ms (memory; memory capacity MC = Σ R² over lags) and the running mean of u over the last 10–500 ms (integration). The same input and noise streams were used in all conditions, with eight independent network replicates per condition. Conditions were: young and older regional parameters; young parameters with only G_srs_ aged; young parameters with only α aged; and G_srs_ scaled by 0.25, 0.5, 1.5 and 2.0 as controls. Paired differences are reported with t-based 95% confidence intervals across replicates.

##### Supplementary Results

###### Nested model comparison

Supplementary Fig. 14 and Supplementary Table 9 report the comparison of four nested models. Median in-sample R² was 0.691, 0.758, 0.745 and 0.985 for M0, M1, M2 and M3, and 95%, 91%, 92% and 0.8% of fits had R² < 0.9 (Supplementary Fig. 14b). M3 had the lowest BIC in 1159 of 1160 participants (Supplementary Fig. 14c). The relay loop alone (M1) improved BIC over M0 in 51% of participants and the reticular pathway (M2) did not improve on M1 (M2 better than M1 in 2%), so the gain came almost entirely from the intrathalamic loop. Blocked cross-validation gave median held-out RMSE of 0.328, 0.309, 0.315 and 0.141 log_10_ units (Supplementary Fig. 14h); the full model was better than M0, M1 and M2 in 98%, 96% and 97% of participants, respectively (paired Wilcoxon, all P = 2.2 × 10^−49^ or smaller).

###### Spectral features

Alpha peaks were detectable in 15%, 77%, 77% and 91% of model spectra (MEG, 92%), and valid knee estimates in 15%, 20%, 13% and 89%. Across participants with a valid estimate in both, model and MEG timescales were correlated at ρ = −0.15, −0.11, −0.23 and 0.87 (Supplementary Fig. 14e shows M0 and M3), and model and MEG alpha frequencies at ρ = 0.02, 0.39, 0.42 and 0.83. The age association of model τ was 0.13, 0.02, 0.09 and −0.33, against −0.42 in MEG (Supplementary Fig. 14f); the age association of model alpha frequency was −0.07 for M3 against −0.17 in MEG, so the full model reproduced the direction of alpha slowing but attenuated it (Supplementary Fig. 14g). The full model also attenuated the age association of τ (−0.33 versus −0.42), so it captures most, not all, of the individual spectral variation.

###### Parameters of the reduced models

In M0, α was at a bound in 100% of fits and β and EMG_a_ in 85% and 78%, indicating that the optimizer was driven to the edge of the allowed region in an attempt to compensate for missing structure. M0 attributed the age effect to cortical gains (G_ee_, ρ = 0.44; G_ei_, ρ = −0.49), but because the fit is poor (median R² 0.69) these associations should not be interpreted as evidence for a cortical mechanism. In particular, the G_ee_ and G_ei_ associations in M0 differ in sign and size from those in M3, which illustrates that parameter–age associations are conditional on the model that is fitted.

###### Fit quality

The corticothalamic model reproduced the cortex-average spectra with a median log-fit R² of 0.985 (Cam-CAN 0.981, OMEGA 0.986, PANS 0.992); 5.1% of fits had R² < 0.95 (minimum 0.887). Fit quality declined with age (Spearman ρ = −0.49), partly reflecting the cohort composition, since the youngest participants were fitted best. Residuals were systematic around 20–25 Hz and near 2 Hz (|mean residual| ≈ 0.05–0.1 log_10_ units). An alpha peak was detectable in 83–95% of the fitted spectra per age bin, compared with 86–96% of the observed spectra.

###### Parameter identifiability

The Jacobian was strongly anisotropic (condition number 4 × 10⁴); the two least constrained directions corresponded to a trade-off between G_ee_ and G_ei_ (loadings −0.73 and +0.64) and to β alone (loading 0.97). Fits reached a bound frequently for β (92% at the upper bound), G_esre_ (77% at the upper bound), α (38% at the upper bound), G_ee_ (18% at the upper bound) and t_0_ (17% at the lower bound). Across independent libraries, parameters estimated from identical-quality fits (χ² ratio 1.001) were reproducible for G_srs_ (ρ = 0.99), α (0.96), t_0_ (0.90) and G_esre_ (0.82; reflecting mainly its position at the bound), and less so for G_ese_ (0.60), EMG_a_ (0.23), G_ei_ (0.15), G_ee_ (0.10) and β (0.08). In synthetic recovery with empirical noise levels, G_srs_ (ρ = 0.91), t_0_ (0.90), α (0.78) and G_ese_ (0.72) were recovered, whereas G_ee_ (0.46), G_ei_ (0.38), G_esre_ (0.37), EMG_a_ (0.48) and β (0.28) were not. Large parameter errors were accompanied by small spectral errors (RMS 0.01–0.05 log_10_ units), indicating that these parameters are partly degenerate. We therefore restrict interpretation to G_srs_, t_0_, α and G_ese_, and to the composite loop gains.

###### Parameters across the lifespan

G_srs_ became more negative with age (pooled ρ = −0.48, 95% CI −0.52 to −0.43), α decreased (−0.33 [−0.38, −0.28]), and G_ese_ (+0.20), G_esre_ (+0.16) and t_0_ (+0.09 [0.03, 0.15]) increased (all q < 0.01). G_ee_, G_ei_, β and EMG_a_ showed no age association (q > 0.3). Among the composite gains, Z increased (+0.42), Y increased (+0.15) and X decreased (−0.11). Because α, t_0_ and several other parameters frequently reached their bounds, age effects on these parameters are likely conservative and were not interpreted quantitatively.

###### Time-domain network

The time-domain network reproduced the analytic spectra (r of log power = 0.997–0.998). Single-unit time constants did not exceed 10.7 ms in the young-adult network or 12.2 ms in the 75-y network, whereas the linearised network contained a non-oscillatory mode with a time constant of 107 ms (102 ms for the 75-y parameters; 88 ms when only G_srs_ was aged) and an alpha-band mode at 9.3 Hz (τ = 53 ms). Changing the unit time constants altered the spectral timescale without affecting the slow network mode or the alpha frequency, whereas changing G_srs_ shifted the slow mode and the spectral timescale together (Spearman ρ = 1.0) and changing t_0_ shifted the alpha frequency. Numerical discretisation at Δt = 0.5 ms underestimated the slow mode by 1.7% and overestimated the alpha-mode frequency by 1.8% relative to Δt = 0.125 ms, with a consistent ordering of young and older networks at all step sizes. With region-specific parameters, the model reproduced the empirical regional timescale map (ρ = 0.95, P_spin_ = 0.001), whereas identical modules did not (ρ = −0.14); cortico-cortical coupling changed the median timescale by only 0.35%.

###### Memory and integration

Across the eight configurations, input-memory capacity decreased with the spontaneous knee timescale (Spearman ρ = −0.98). Relative to the young network, MC was higher in the older network (+0.13, 95% CI 0.09–0.17) and when only G_srs_ was aged (+0.33, 0.29–0.36), and essentially unchanged when only α was aged (−0.02, −0.03 to 0.00). The memory kernel peaked at a lag of ∼50 ms, consistent with the thalamocortical loop delay. Thus, within this network, a shorter spontaneous timescale was not accompanied by reduced input memory.

###### Knee frequency relative to the fitted band

The validity criterion (0 < τ < 500 ms) corresponds to *f*_k_ > 0.32 Hz and therefore does not by itself place the knee inside the 2–45 Hz fitting band. Of 75,670 valid regional fits (from 78,608 spectra of participants with age), 64,357 (95.0%) had a knee between 2 and 45 Hz, 3,394 (5.0%) below 2 Hz and 8 above 45 Hz. The share inside the band was 98.3% in Cam-CAN, 96.0% in OMEGA and 86.2% in PANS, and 82.6%, 96.5% and 98.0% in participants younger than 18 years, aged 18–59 years and aged 60 years or older. Knees below 2 Hz are long timescales (τ > 80 ms) that were retained in the primary analysis and were concentrated in children, in whom the knee often lies at or below the lower edge of the band; this is also where most invalid fits occurred (Supplementary Fig. 2). When each participant’s median was restricted to in-band knees, the age association was ρ = −0.53 (95% CI −0.57 to −0.49; n = 1,155), compared with −0.50 in the primary analysis, and ρ = −0.51 when at least 34 in-band regions were also required (n = 989). The association therefore did not depend on knees extrapolated beyond the fitted band.

###### Alignment with the functional gradient across life stages

The correspondence between the regional τ map and the principal functional gradient was ρ = 0.53 in youth, 0.57 in adults and 0.37 in older adults. For the differences, spin-permutation P values were 0.32 (adults minus older adults, Δρ = 0.20), 0.53 (youth minus older adults, Δρ = 0.16) and 0.66 (youth minus adults, Δρ = −0.04). Participant-level bootstrap 95% intervals (300 resamples within cohort and life stage) were 0.07 to 0.30, 0.02 to 0.24 and −0.11 to 0.01. The spin test accounts for spatial autocorrelation but not for sampling error in the maps, and the bootstrap does the opposite, so the two answer different questions; together they support at most a suggestive weakening of alignment in older adults.

###### Age change in the fitted spectra

Between participants aged 20–29 years (n = 188) and 70–89 years (n = 180), the median τ of the MEG spectra fell by 45% and that of the full-model spectra by 38%, and the median alpha frequency fell by 1.31 Hz in the MEG spectra and 0.73 Hz in the full-model spectra (group medians pooled over cohorts, not adjusted for cohort). The fitted parameters therefore capture most of the shortening of τ but about half of alpha slowing. In the independent-substitution analysis (Supplementary Table 6), the age-related decrease in α raised the model alpha peak by 0.21 Hz and offset the −0.18 Hz effect of t_0_, so that shifting all parameters together gave +0.17 Hz; the substitution analysis ignores correlations among parameters and should not be read as a prediction of the alpha trajectory.

#### Limitations

Participant-level corticothalamic parameters were estimated from cortex-average spectra; regional parameters come from a 280-participant subsample and from group-level fits. Several parameters (G_ee_, G_ei_, β, EMG_a_, G_esre_) were not individually identifiable, and parameters at their bounds limit inference on α and t_0_; conclusions are therefore restricted to G_srs_, t_0_, α and G_ese_. Identifiability and recovery analyses used spectra generated by the model itself and therefore represent optimistic bounds. Residuals showed systematic misfit at 20–25 Hz. The network simulations used n = 8 replicate networks, a fixed readout based on cortical activity, and random input weights; they illustrate model implications and were not validated against behaviour. Circuit architectures other than the nested versions of this field model (Supplementary Fig. 14) and the posterior uncertainty of the parameters were not evaluated.

#### Supplementary Figures

**Supplementary Fig. 1.**
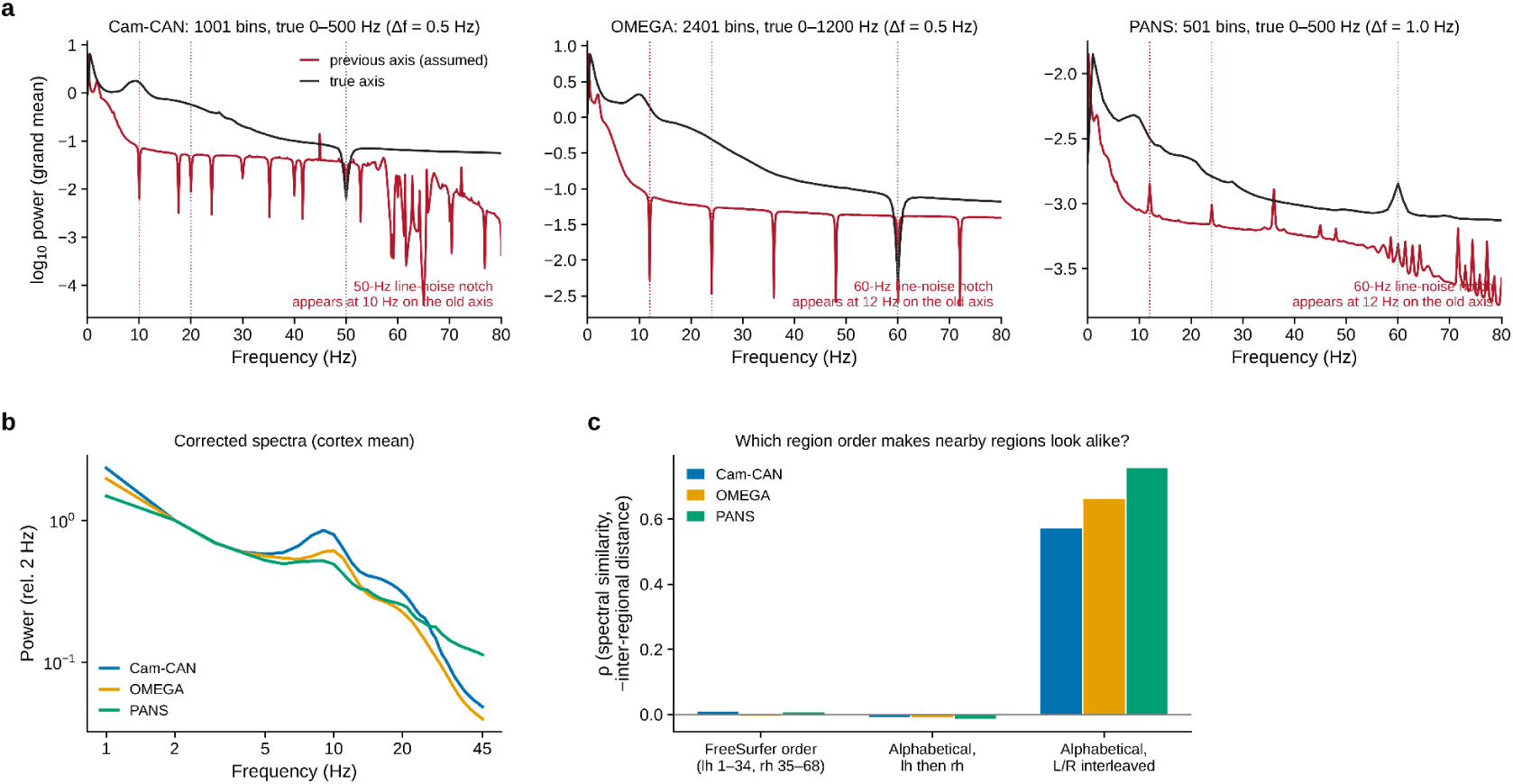
| Frequency-axis correction and region-order verification. **a,** Cortex-mean (grand-mean) log₁₀ power spectra for Cam-CAN (1,001 bins; true range 0–500 Hz, Δf = 0.5 Hz), OMEGA (2,401 bins; 0–1,200 Hz, Δf = 0.5 Hz) and PANS (501 bins; 0–500 Hz, Δf = 1.0 Hz). Red, spectra plotted on the previously assumed frequency axis; black, the same spectra on the true axis. On the assumed axis, the line-noise notch and its harmonics appeared at the wrong frequency (50 Hz appeared at 10 Hz in Cam-CAN; 60 Hz appeared at 12 Hz in OMEGA and PANS; dotted vertical lines). On the true axis, the notch (or peak, in PANS) falls at 50 Hz (Cam-CAN) or 60 Hz (OMEGA, PANS). **b,** Corrected cortex-mean spectra (1–45 Hz, the analysis range), normalised to the power at 2 Hz and plotted on log–log axes (blue, Cam-CAN; orange, OMEGA; green, PANS). **c,** Test of the region order. For each cohort, Spearman’s ρ between spectral similarity and negative inter-regional distance is shown for three candidate orderings of the 68 Desikan–Killiany regions (FreeSurfer order, lh 1–34 then rh 35–68; alphabetical, lh then rh; alphabetical, left/right interleaved). Only the interleaved ordering makes neighbouring regions spectrally similar (ρ ≈ 0.57–0.75), and it was therefore used throughout.

**Supplementary Fig. 2.**
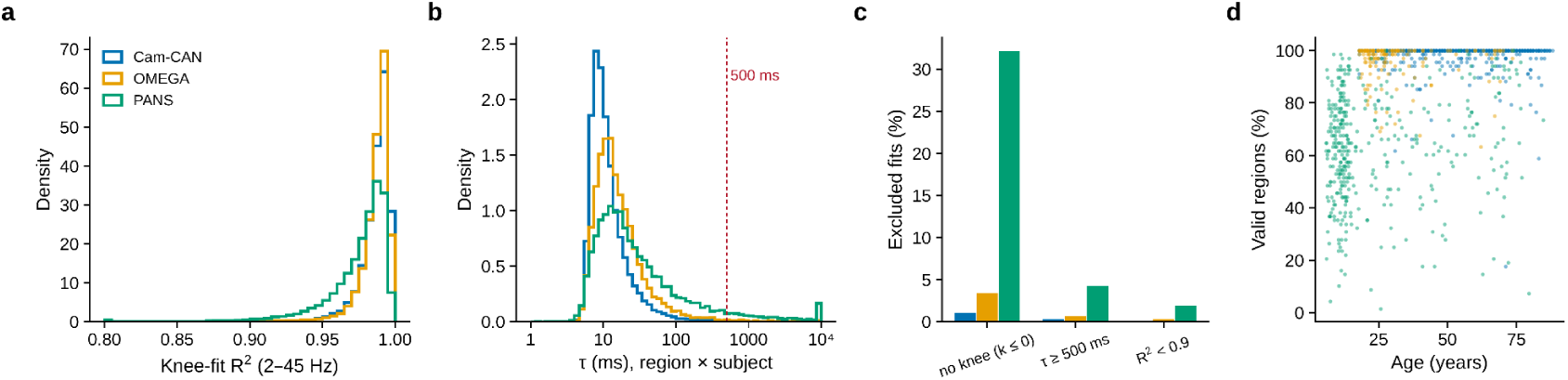
| Quality control of the knee fits. **a,** Distribution of the goodness of fit (R², 2–45 Hz) of the aperiodic-with-knee model across all region × participant fits, shown as density histograms for each cohort (blue, Cam-CAN; orange, OMEGA; green, PANS). **b,** Distribution of the estimated τ = 1/(2π f_k_) for all region × participant fits (log-scaled x-axis). The red dashed line marks the 500 ms upper limit for a fit to count as valid; values beyond the plotted range are collected in the last bin. **c,** Percentage of fits excluded in each cohort, by reason: no knee (knee parameter k ≤ 0), τ ≥ 500 ms, or R² < 0.9. A fit was valid only if none of these applied. **d,** Percentage of the 68 regions with a valid fit per participant, against age. Each dot is one participant, coloured by cohort. Valid-region coverage was highest in Cam-CAN and lowest in the youngest PANS participants, which is why the participant-level τ is the median over valid regions and why we tested sensitivity to the minimum number of valid regions (Supplementary Fig. 5).

**Supplementary Fig. 3.**
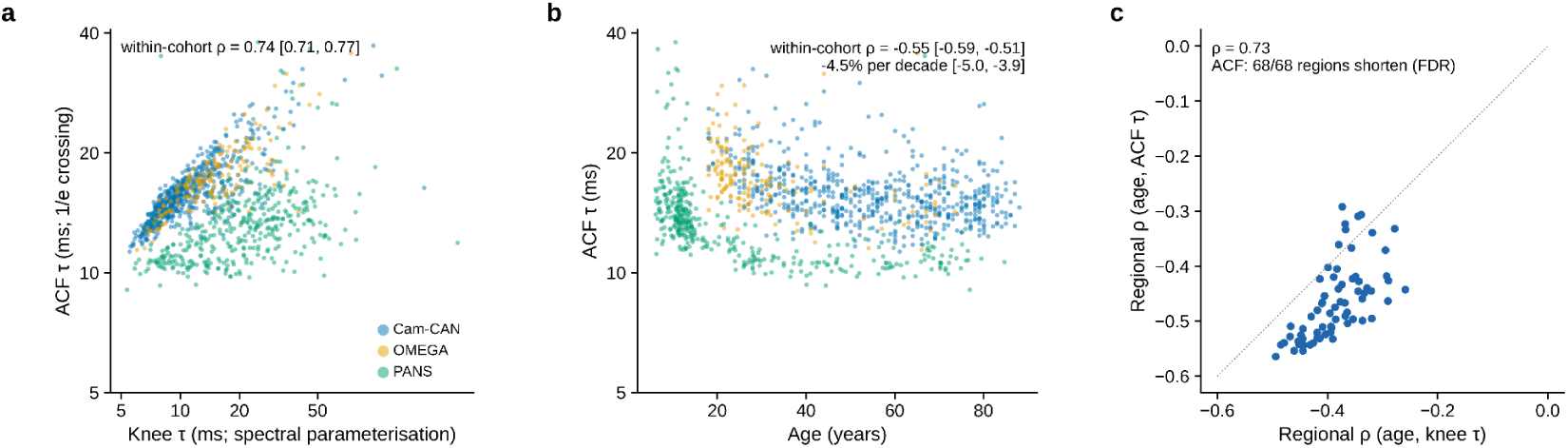
| Autocorrelation-based timescale. a, Autocorrelation τ (1/e crossing) against knee τ. b, Autocorrelation τ against age. c, Regional age effects of autocorrelation τ against those of knee τ.

**Supplementary Fig. 4.**
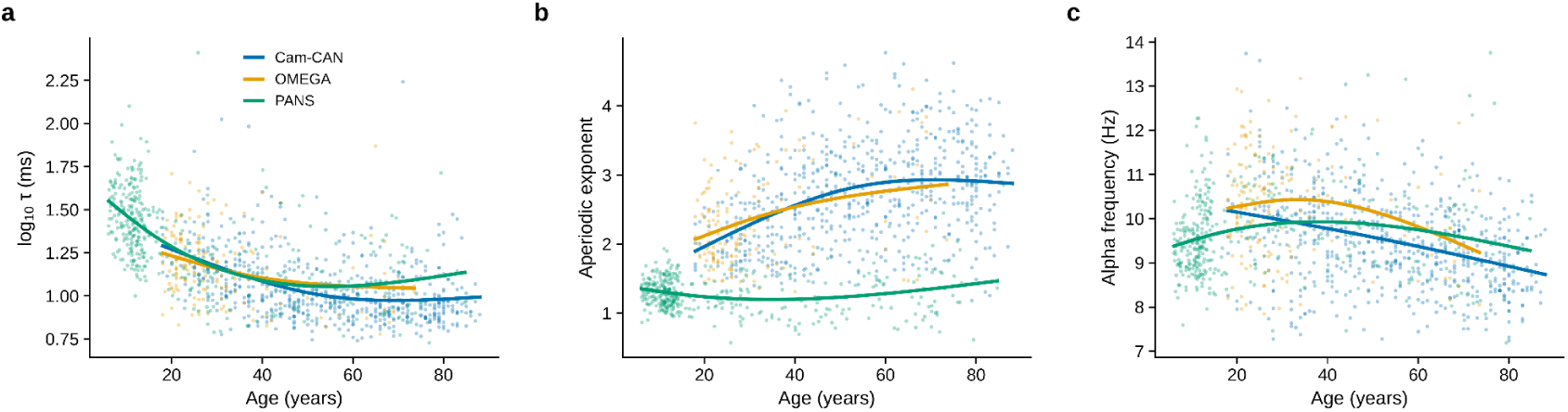
| Cohort-specific lifespan trajectories of the spectral features. a, log₁₀ τ; b, aperiodic exponent; c, alpha peak frequency, against age for Cam-CAN (blue, n = 606), OMEGA (orange, n = 162) and PANS (green, n = 388; n = 387 for alpha frequency). Dots, participants; lines, cohort-specific natural cubic splines (3 d.f.). Within-cohort Spearman correlations with age: log₁₀ τ, Cam-CAN ρ = −0.434 (P = 3.436 × 10⁻²⁹), OMEGA ρ = −0.333 (P = 1.464 × 10⁻⁵), PANS ρ = −0.640 (P = 4.875 × 10⁻⁴⁶), meta-analysis ρ = −0.498 [−0.540, −0.453] (P = 1.716 × 10⁻⁷⁶); aperiodic exponent, ρ = 0.364 (P = 1.785 × 10⁻²⁰), 0.250 (P = 0.001), −0.113 (P = 0.026), meta ρ = 0.196 [0.139, 0.251] (P = 1.911 × 10⁻¹¹); alpha frequency, ρ = −0.358 (P = 9.768 × 10⁻²⁰), −0.112 (P = 0.157), 0.132 (P = 0.009), meta ρ = −0.166 [−0.222, −0.110] (P = 1.311 × 10⁻⁸). Full statistics in Supplementary Table 3a,b. The PANS exponent is lower overall than in the other cohorts (different system and 1-Hz resolution); age effects are therefore estimated within cohort.

**Supplementary Fig. 5.**
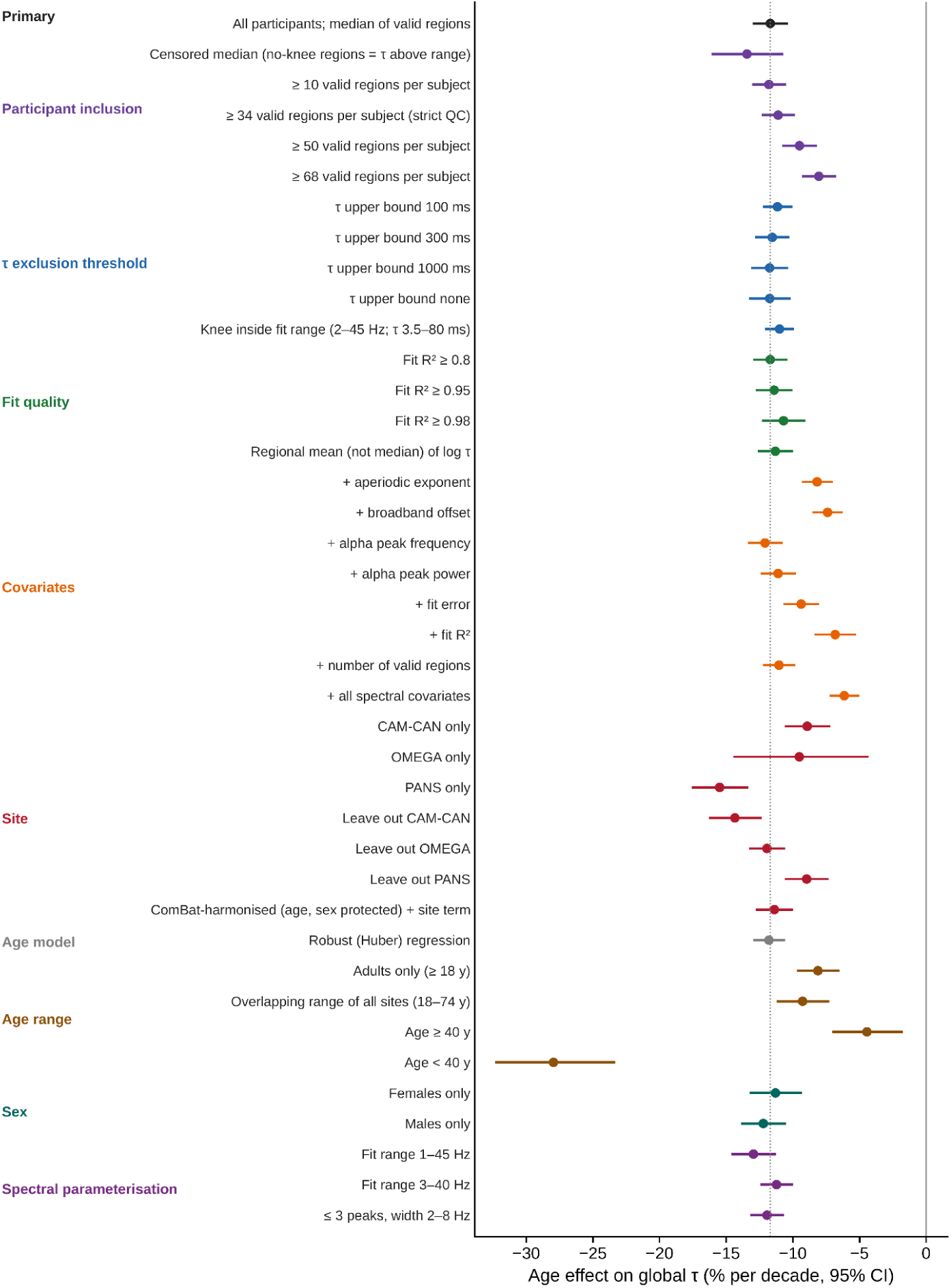
| Robustness of the age association with global intrinsic timescales. Forest plot showing the estimated percentage change in global intrinsic timescales per decade across 40 alternative analytical specifications. Points indicate effect estimates and horizontal bars denote 95% confidence intervals. The primary analysis is highlighted in black, and the vertical dashed line indicates its estimated effect. Specifications varied participant inclusion criteria, spectral fitting parameters, quality-control thresholds, covariate adjustment, cohort handling, age ranges and sex stratification. All specifications yielded negative age associations, although effect magnitudes differed across analytical choices. Full numerical results are provided in Supplementary Table 2.

**Supplementary Fig. 6.**
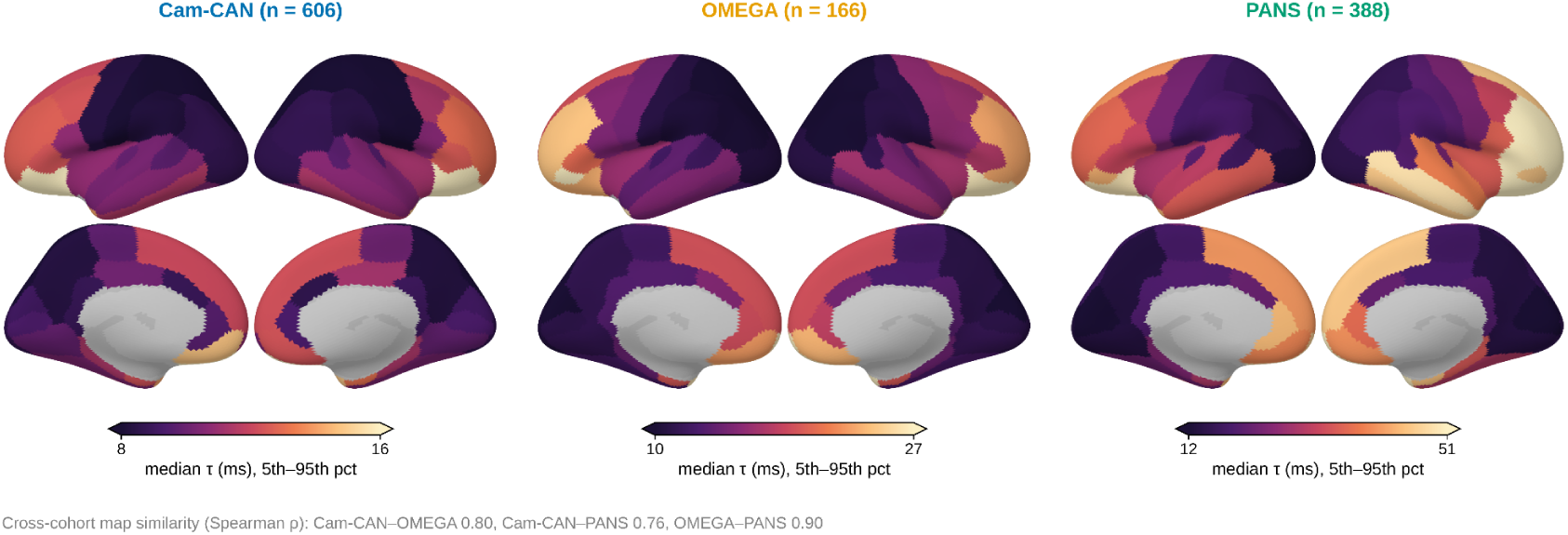
| Regional τ maps in the three cohorts. Cortical maps of median regional τ (ms) in Cam-CAN (n = 606), OMEGA (n = 162) and PANS (n = 388), each shown in lateral (top) and medial (bottom) views of both hemispheres. The colour scale of each map is set to the 5th–95th percentile of that cohort’s regional medians (Cam-CAN 8–16 ms; OMEGA 10–27 ms; PANS 12–51 ms), so absolute values should be compared with the colour bars, and the spatial pattern across maps. In all cohorts τ is shortest in sensorimotor and posterior regions and longest in frontal and temporal association regions. Cross-cohort spatial similarity (Spearman ρ across 68 regions) is 0.80 (Cam-CAN–OMEGA), 0.76 (Cam-CAN–PANS) and 0.90 (OMEGA–PANS).

**Supplementary Fig. 7.**
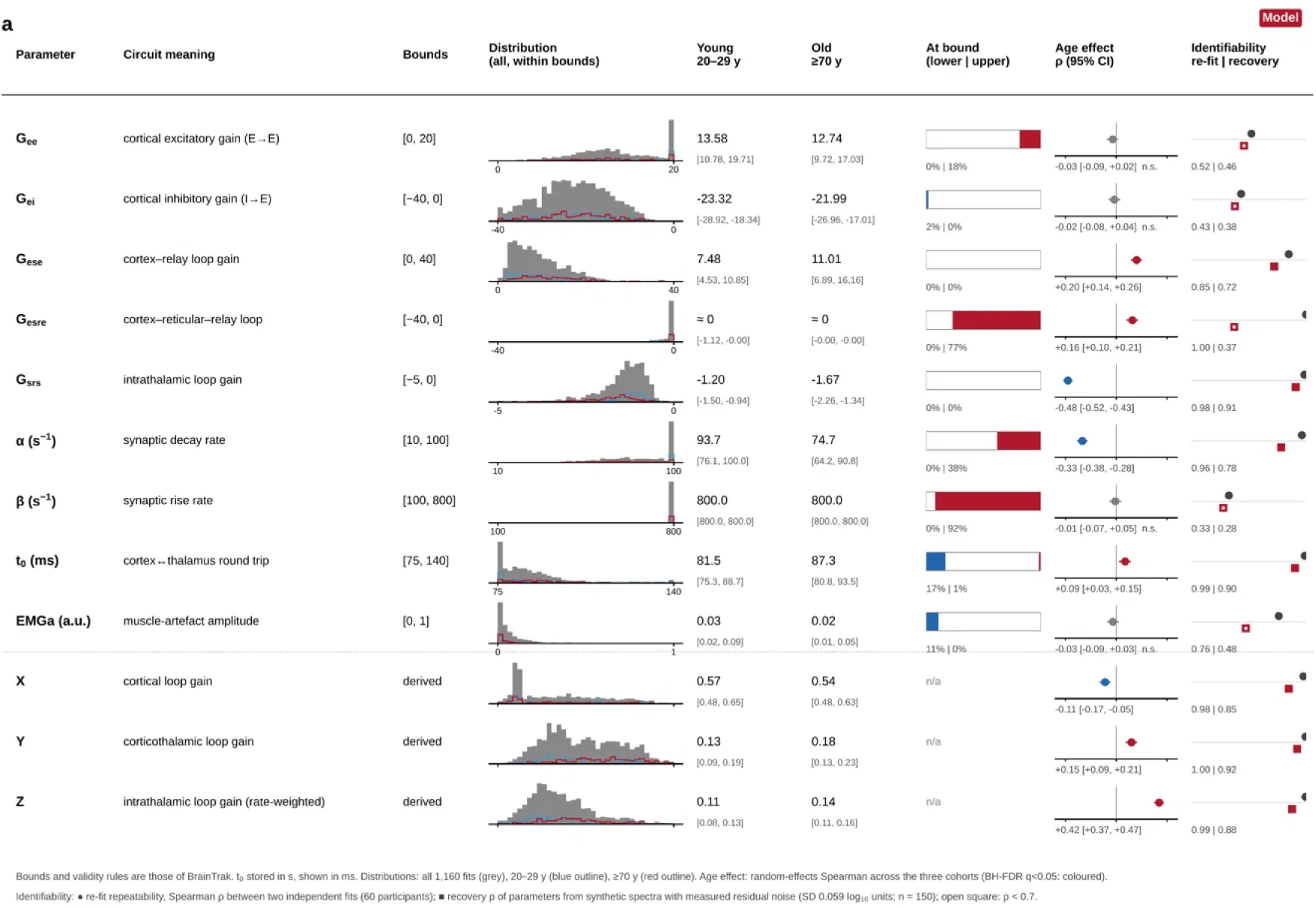
| Parameter dictionary of the corticothalamic model. For each fitted parameter (G_ee_, G_ei_, G_ese_, G_esre_, G_srs_, α, β, t_0_, EMG_a_) and each composite loop gain (X, Y, Z), the panel lists its circuit meaning, its bounds, and its distribution across all 1,160 fits (grey; blue outline, 20–29 y, n = 194; red outline, ≥70 y, n = 180). It also gives the median and interquartile range in the two age groups. The “At bound” column shows the fraction of fits at the lower (blue) and upper (red) bound, with a tolerance of 1% of the parameter range. The “Age effect” column shows the Spearman correlation with age, pooled across the three cohorts by inverse-variance weighting on the Fisher z scale. The point is the pooled ρ and the bar is the 95% confidence interval; coloured points have Benjamini–Hochberg q < 0.05, red positive and blue negative, and grey points are not significant. The “Identifiability” column gives the Spearman ρ between independent re-fits (circle, n = 60) and the recovery of the parameter from synthetic spectra containing empirical noise (square, n = 150; open if ρ < 0.7). t_0_ is stored in seconds and shown in milliseconds. Several parameters lie at a bound in many fits (β 92%, G_esre_ 77%, α 38%, G_ee_ 18%, t_0_ 17% at the lower bound). Only G_srs_, t_0_, α and G_ese_ were reliably recovered.

**Supplementary Fig. 8.**
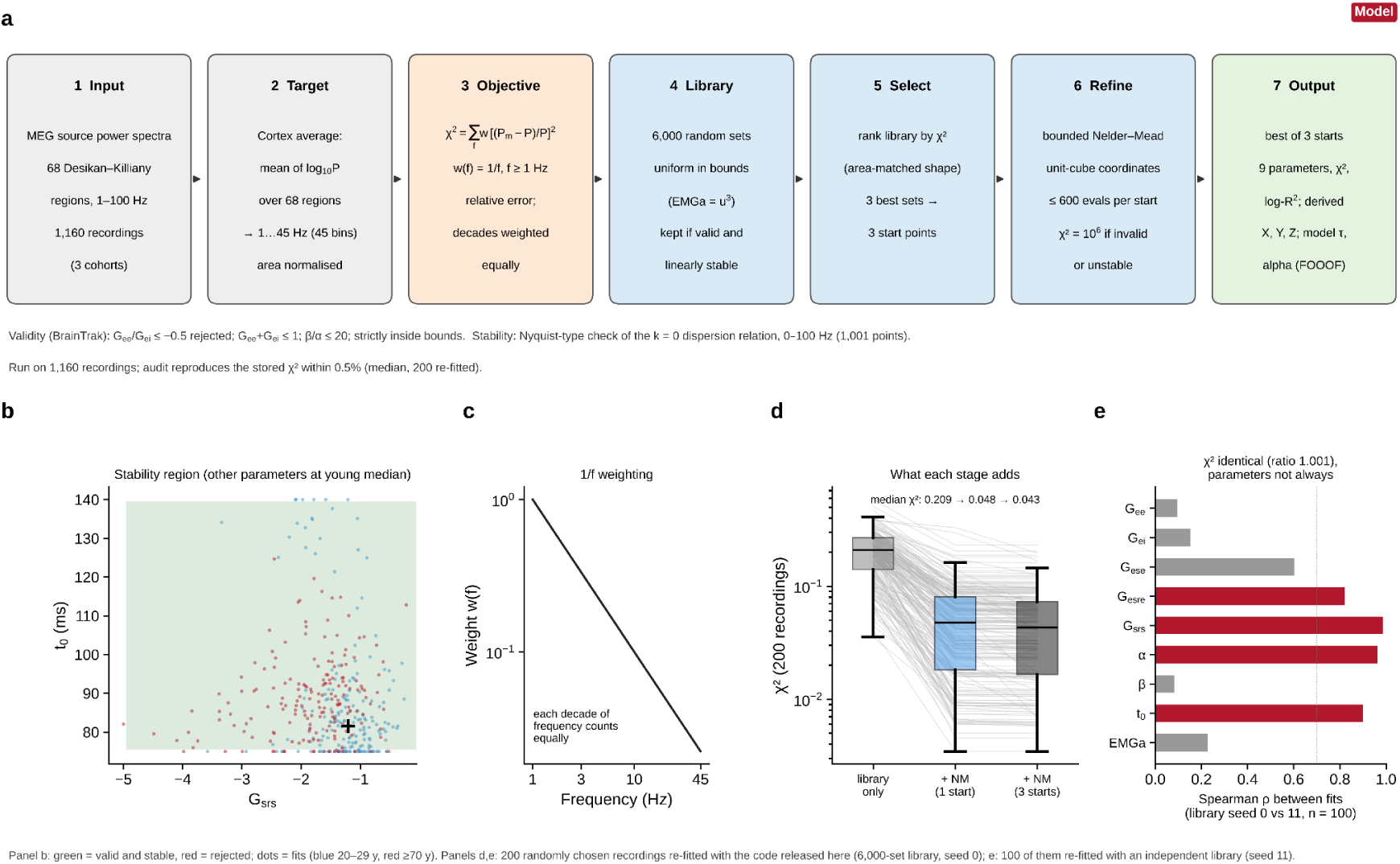
| Fitting procedure for the corticothalamic neural-field model. **a**, Pipeline. The target is the cortex-average spectrum: the mean of log_10_ power over the 68 Desikan–Killiany regions, restricted to 1–45 Hz (45 bins) and area-normalised. The objective is χ² = Σf w(f)[(Pm − P)/P]² with w(f) = 1/f. A library of 6,000 valid and linearly stable random parameter sets is ranked by χ². The three best sets start a bounded Nelder–Mead refinement (≤ 600 evaluations per start, unit-cube coordinates; invalid or unstable points receive χ² = 10⁶). The best solution is kept, together with log-fit R², composite gains X, Y, Z and the model τ and alpha peak (FOOOF). Validity requires G_ee_/G_ei_ > −0.5, G_ee_ + G_ei_ ≤ 1, β/α ≤ 20 and parameters strictly inside their bounds. Stability is assessed with a Nyquist-type criterion on the k = 0 dispersion relation (0–100 Hz, 1,001 points). **b**, Region of the G_srs_–t_0_ plane that is valid and stable (green) with the other parameters at the young-adult median; rejected sets would appear in red. Dots, fitted values (blue, 20–29 y; red, ≥70 y). **c**, The 1/f weighting gives each decade of frequency approximately equal weight. **d**, χ² of 200 randomly selected recordings after the library stage alone, after refinement from one start, and from three starts (grey lines, individual recordings; boxes, median and interquartile range). Median χ² is 0.209, 0.048 and 0.043, respectively. **e**, Spearman correlation between parameters estimated with two independently drawn libraries (seeds 0 and 11, n = 100). χ² values were nearly identical (median ratio 1.001), but several parameters were not reproduced (grey bars, ρ ≤ 0.7; red bars, ρ > 0.7).

**Supplementary Fig. 9.**
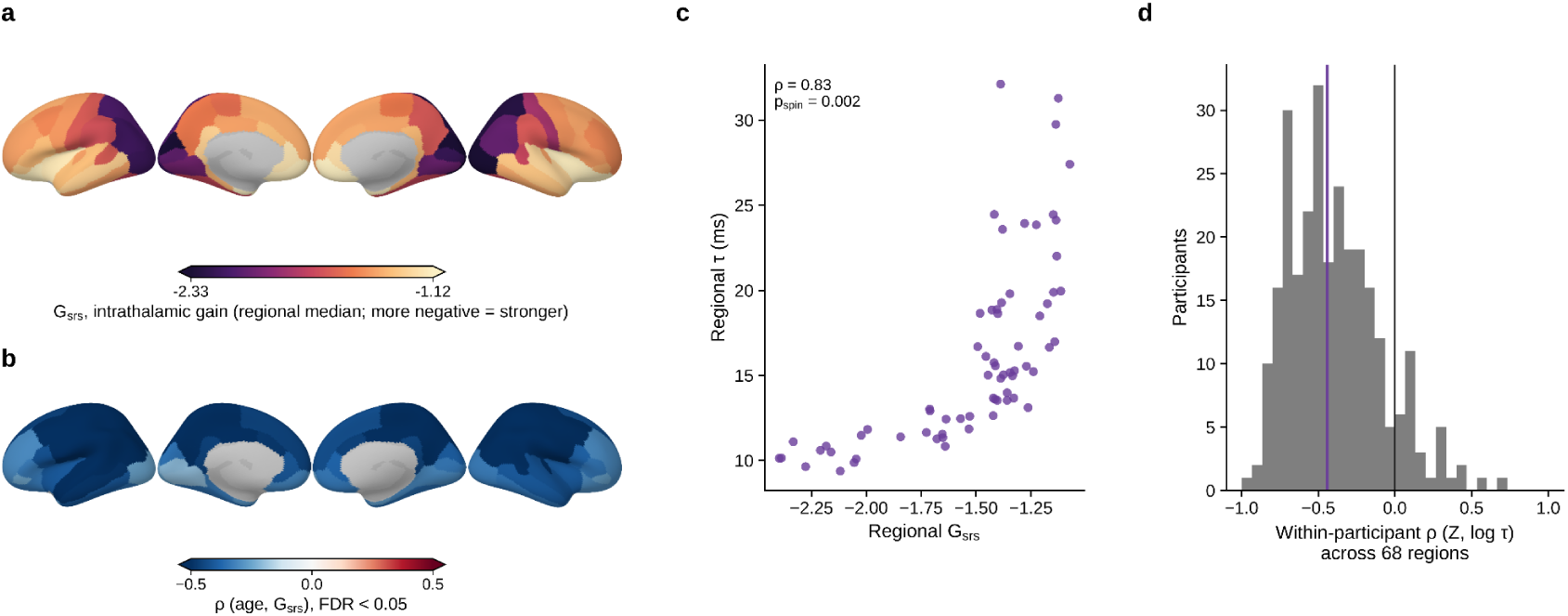
| The intrathalamic loop gain G_srs_, regional τ and age. **a,** Regional median G_srs_ (intrathalamic loop gain; more negative values indicate a stronger loop) across participants (regional fits in 280 participants), shown on the cortical surface (lateral and medial views of both hemispheres). **b,** Regional associations between age and intrathalamic feedback gain (G_srs_), quantified using Spearman correlations. Negative correlations indicate that G_srs_ becomes more negative with age, corresponding to stronger effective intrathalamic feedback. Blue, ρ < 0 (G_srs_ becomes more negative, that is, the intrathalamic loop strengthens, with age); red, ρ > 0. Only regions surviving false discovery rate correction (FDR < 0.05) are coloured; all 68 regions showed a significant negative association. **c,** Regional median τ against regional median G_srs_ across the 68 regions (Spearman ρ = 0.83; spin-test p = 0.002). **d,** Within-participant Spearman’s ρ between the model-derived intrathalamic term Z and log τ across the 68 regions, as a histogram over participants. The black line marks ρ = 0, and the purple line marks the median (about −0.45), showing that the spatial relation in **c** holds within individuals and not only in the group average.

**Supplementary Fig. 10.**
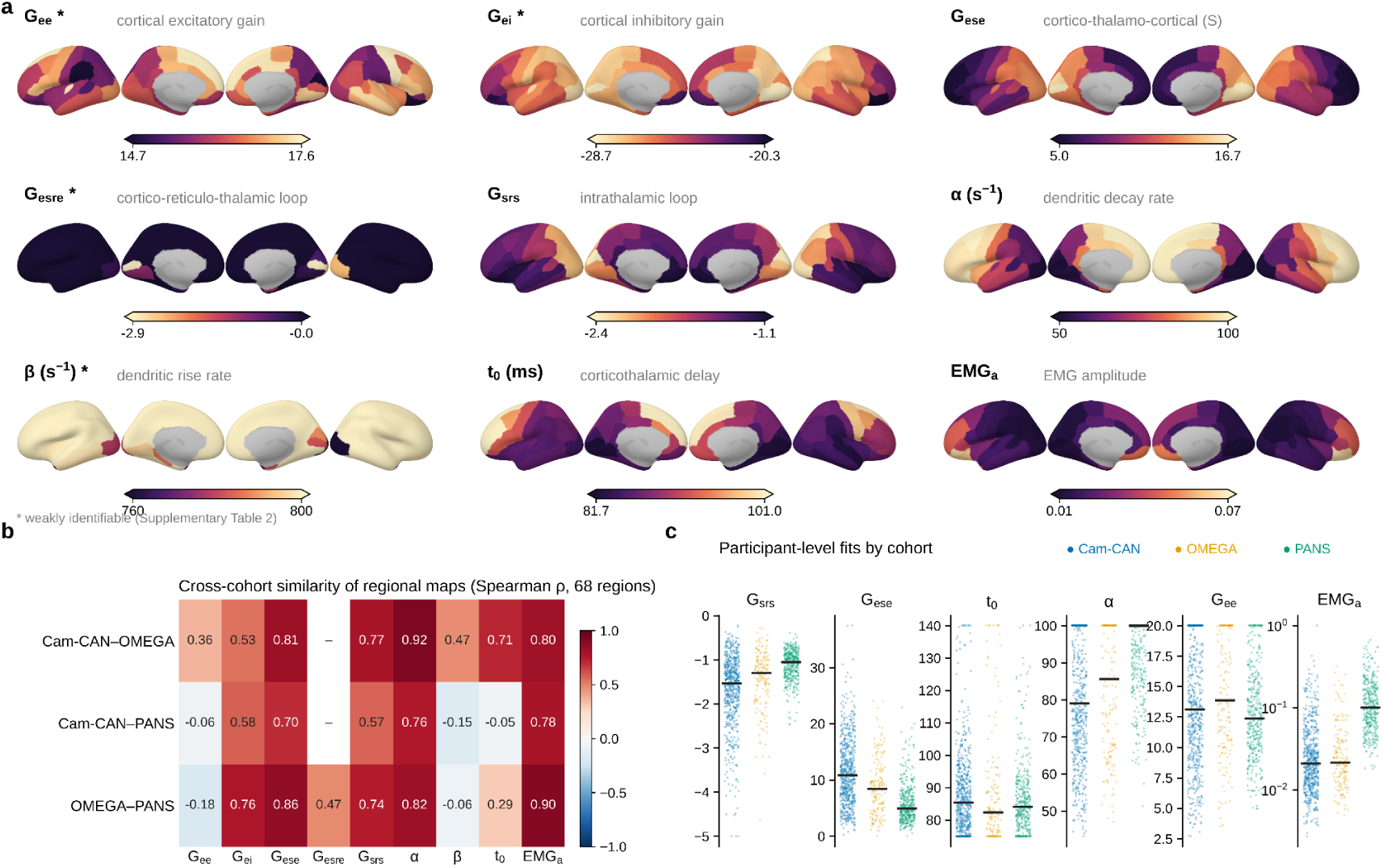
| Regional maps of the corticothalamic parameters and their reproducibility across cohorts. **a,** Cortical maps of regional median parameters of the corticothalamic model: G_ee_ (cortical excitatory gain), G_ei_ (cortical inhibitory gain), G_ese_ (cortico-thalamo-cortical loop), G_esre_ (cortico-reticulo-thalamic loop), G_srs_ (intrathalamic loop), α and β (dendritic decay and rise rates, s⁻¹), t₀ (corticothalamic delay, ms) and EMG_a_ (EMG amplitude). Colour ranges are shown under each map (note the narrow ranges for G_ee_, G_ei_ and β, and that G_esre_ is near zero in most regions). An asterisk marks parameters that are weakly identifiable in simulation (Supplementary Table 4); their maps should be read with caution. **b,** Cross-cohort similarity of the regional maps (Spearman ρ across 68 regions) for each pair of cohorts and each parameter; “–” indicates that the map could not be compared (G_esre_, Cam-CAN, owing to almost no variation across regions). Maps of G_srs_, α, G_ese_ and EMG_a_ replicate across all three pairs (ρ ≥ 0.57), whereas G_ee_, β and t₀ do not consistently replicate. **c,** Participant-level fitted values by cohort for six parameters (dots, participants; horizontal black lines, medians; EMG_a_ on a log axis). Piles of values at the limits of the bounds (e.g. α = 100 s⁻¹, t₀ = 75 and 140 ms, G_ee_ = 20) mark where the optimiser reached its constraint.

**Supplementary Fig. 11.**
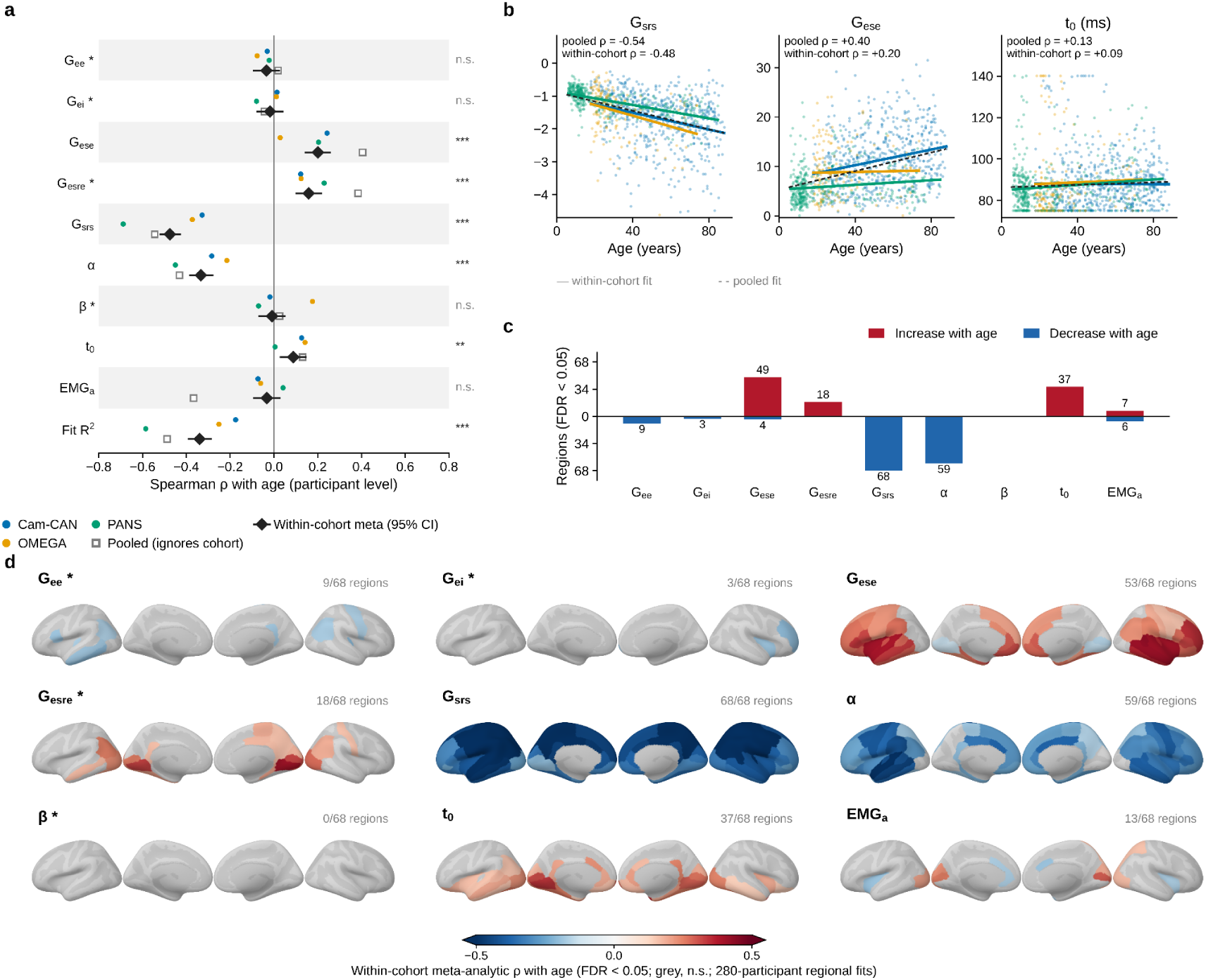
| Age effects on the corticothalamic parameters. Panels a and b show the cortex-average fits of all 1,156 participants with age; panels c and d show the regional fits of 280 participants. **a,** Spearman’s ρ between age and each parameter from the cortex-average fit of each participant (n = 1,156) and the goodness of fit (Fit R²). Coloured dots, individual cohorts (blue, Cam-CAN; orange, OMEGA; green, PANS); black diamonds, the within-cohort meta-analytic estimate with 95% CI (Fisher-z, inverse-variance weights); open squares, the pooled estimate that ignores cohort. Right-hand labels give the significance of the meta-analytic effect after BH-FDR correction (n.s., not significant; ** and *** by FDR level). Asterisks after parameter names mark weakly identifiable parameters. **b,** Participant-level G_srs_, G_ese_ and t₀ against age. Solid lines, within-cohort fits; dashed black line, pooled fit; each panel gives the pooled and the within-cohort ρ (G_srs_ −0.54 and −0.48; G_ese_ +0.40 and +0.20; t₀ +0.13 and +0.09). The pooled estimates are larger than the within-cohort ones because cohorts differ in age range and in level, which is why the within-cohort estimate is the primary one. **c,** Number of regions with a significant age association (FDR < 0.05; regional fits in 280 participants) for each parameter, split into increases (red) and decreases (blue) with age: G_srs_ decreases in all 68 regions, α in 59, and G_ese_ increases in 49 (and decreases in 4); β shows none. **d,** Regional maps of the within-cohort meta-analytic ρ with age for each parameter (FDR < 0.05; grey, not significant; regional fits in 280 participants). The number of significant regions is given above each map.

**Supplementary Fig. 12.**
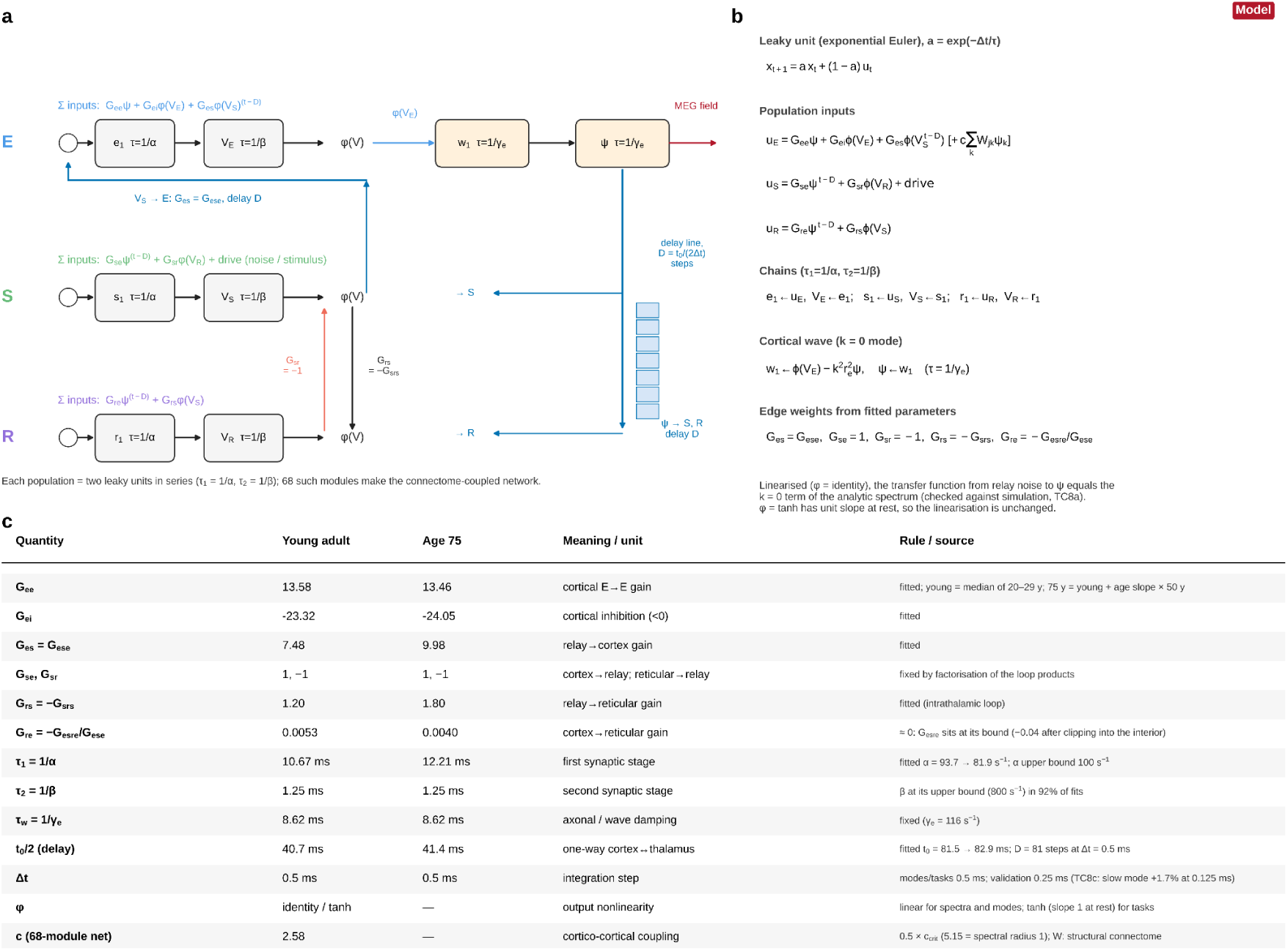
| Architecture, update rule and parameters of the thalamocortical rate network. **a**, Each population (cortical E, relay S, reticular R) consists of two leaky units in series with time constants 1/α and 1/β. The cortical field ψ, which corresponds to the MEG-observable activity, is obtained through two further units with time constant 1/γ_e_. Inputs to each population are listed above its row. Delay lines of D = t_0_/(2Δt) steps carry ψ → S, ψ → R and VS → E; the S → R and R → S connections have no delay. Sixty-eight such modules form the connectome-coupled network. **b**, Update rule (exponential Euler integration, a = exp(−Δt/τ)), population inputs, chains, cortical wave equation, and the mapping from fitted parameters to edge weights: G_es_ = G_ese_, G_se_ = 1, G_sr_ = −1, G_rs_ = −G_srs_, G_re_ = −G_esre_/G_ese_. φ is the identity for spectral analysis and tanh for task simulations, and c is the cortico-cortical coupling. With φ equal to the identity, the transfer function from relay noise to the cortical field ψ reproduces the k = 0 component of the corresponding analytical corticothalamic model, as described in the Supplementary Methods. **c**, Parameter values for the young-adult network (median of 20–29 y) and the 75-y network (young value plus the age regression slope × 50 y), with their meaning and source. Single-unit time constants are ≤ 10.7 ms (young adult) and ≤ 12.2 ms (age 75). G_esre_ sits at its bound and was set to −0.04 before conversion to G_re_. β is at its upper bound (800 s⁻¹) in 92% of fits. The integration step is 0.5 ms for modes and tasks.

**Supplementary Fig. 13.**
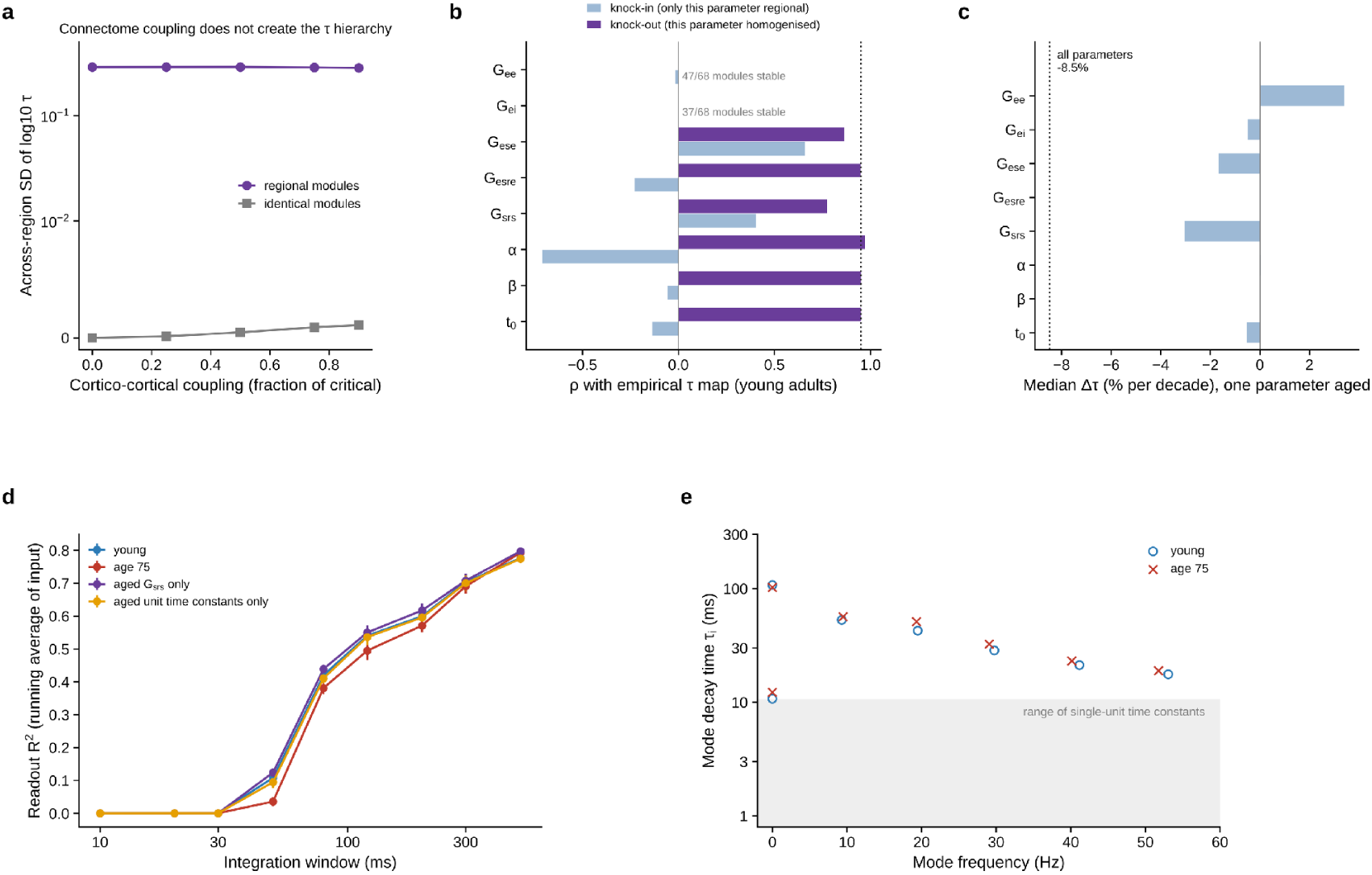
| Connectome thalamocortical RNN. a, Across-region spread of τ as a function of cortico-cortical coupling for regionally fitted versus identical modules. b, Correlation with the empirical young-adult τ map when one parameter is regional (knock-in) or homogenised (knock-out); some knock-ins produce unstable modules. c, Median regional change in τ when only one parameter is aged; dotted line, all parameters aged. d, Temporal integration: readout R² for the running average of the input over windows of 10–500 ms for the young-adult, age-75, aged-intrathalamic-gain-only and aged-unit-time-constants-only networks (mean ± s.d. across eight input streams); performance increases with window duration and is broadly comparable across configurations. e, Full eigenmode spectrum of the linearised single-module network (mode decay time against mode frequency) for young-adult (circles) and age-75 (crosses) parameters; grey band, range of single-unit time constants.

**Supplementary Fig. 14.**
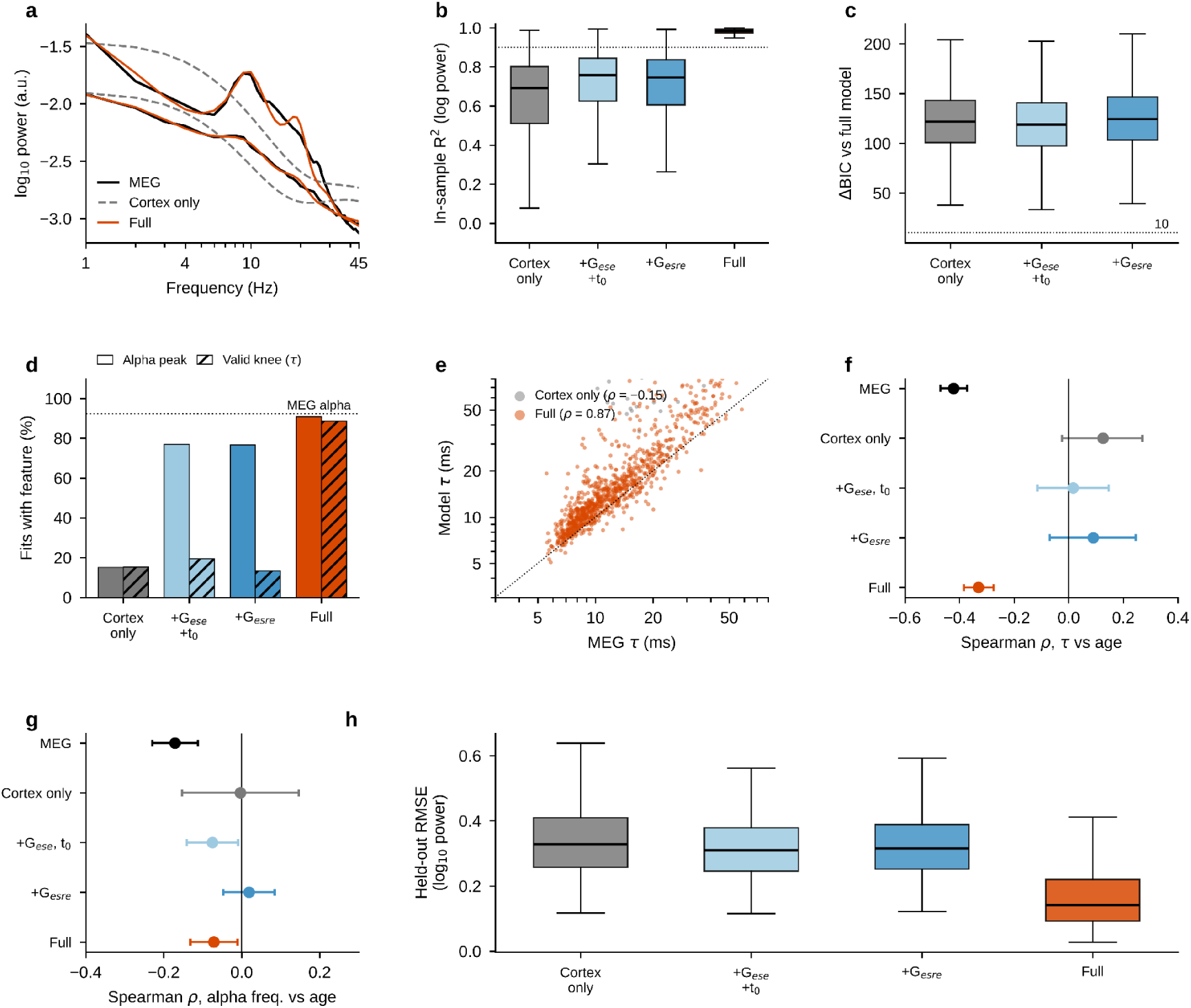
| A cortex-only field does not reproduce the spectra; thalamic terms are required. **a,** Cortex-average MEG spectra of a younger (12 years) and an older (70 years) participant (second spectrum offset for display) with the fits of the cortex-only model (dashed grey) and the full model (orange). **b,** In-sample fit quality (R² of log_10_ power) for the four nested models (box, interquartile range; line, median; whiskers, 1.5 × interquartile range; n = 1,160 participants). The dotted line marks R² = 0.9. **c,** Difference in Bayesian information criterion (ΔBIC) between each reduced model and the full model; positive values favour the full model, and the dotted line marks ΔBIC = 10. **d,** Percentage of model spectra from which an alpha peak (7–14 Hz, open bars) or a valid knee (τ, hatched bars) could be recovered by spectral parameterization; the dotted line marks the percentage of MEG spectra with an alpha peak. **e,** Timescales re-estimated from model spectra for the cortex-only (grey) and full (orange) models against MEG timescales; the dotted line shows equality. **f,g,** Within-cohort pooled Spearman correlations (points) with 95% confidence intervals (bars) between age and the timescale (f) or alpha frequency (g) of MEG spectra and of each model’s spectra. **h,** Held-out error (RMSE of log_10_ power) from blocked cross-validation across five 9-Hz blocks (n = 300 participants). M0, cortex only; M1, + relay loop (G_ese_, t_0_); M2, + reticular pathway (G_esre_); M3, full model.

#### Supplementary Tables

**Supplementary Table 1.** | Cohorts and analysed samples.

| Cohort | MEG system | Rest | Recorded (*n*) | Analysed (*n*) | Age range (y) | Age, mean ± s.d. | Female | Original spectrum | Valid regional fits | Median τ (ms) |
|---|---|---|---|---|---|---|---|---|---|---|
| Cam-CAN | Elekta VectorView, 306 ch, 1 kHz | eyes closed, ~8.5 min | 606 | 606 | 18–88 | 54.2 ± 18.3 | 299 | 0–500 Hz, Δf 0.5 Hz | 98.6% | 10.0 |
| OMEGA | CTF, 275 ch, 2.4 kHz | eyes open, ≥ 5 min | 166 | 162 | 18–74 | 31.6 ± 14.6 | 80 | 0–1,200 Hz, Δf 0.5 Hz | 95.6% | 14.4 |
| PANS | MEGIN, 306 ch, 1 kHz | eyes closed, 5–8 min | 388 | 388 | 6–85 | 25.3 ± 20.9 | 183 | 0–500 Hz, Δf 1 Hz | 92.9% | 21.2 |

*Note.* Valid regional fits, percentage of region × participant knee fits that met the quality criteria (knee > 0, τ < 500 ms, *R*<sup>2</sup> ≥ 0.9).

**Supplementary Table 2.** | Sensitivity of the global age effect on τ.

| Analysis | *n* | % per decade (95% CI) | Meta ρ |
|---|---|---|---|
| *Primary analysis* |  |  |  |
| All participants, median of valid regions (2–45 Hz, τ < 500 ms, *R*<sup>2</sup> ≥ 0.9) | 1,156 | −11.7 (−12.9 to −10.5) | −0.50 |
| *Participant inclusion* |  |  |  |
| Censored median (no-knee regions = τ above range) | 1,156 | −13.4 (−16.0 to −10.8) | −0.38 |
| ≥ 10 valid regions per subject | 1,151 | −11.8 (−13.0 to −10.6) | −0.50 |
| ≥ 34 valid regions per subject (strict QC) | 1,057 | −11.1 (−12.3 to −9.9) | −0.50 |
| ≥ 50 valid regions per subject | 892 | −9.5 (−10.7 to −8.3) | −0.48 |
| ≥ 68 valid regions per subject | 579 | −8.1 (−9.2 to −6.9) | −0.46 |
| *τ exclusion threshold* |  |  |  |
| T upper bound 100 ms | 1,155 | −11.2 (−12.2 to −10.1) | −0.53 |
| T upper bound 300 ms | 1,156 | −11.6 (−12.7 to −10.3) | −0.50 |
| T upper bound 1000 ms | 1,156 | −11.8 (−13.0 to −10.4) | −0.49 |
| T upper bound none | 1,156 | −11.7 (−13.2 to −10.2) | −0.48 |
| Knee inside fit range (2–45 Hz; τ 3.5–80 ms) | 1,155 | −11.0 (−12.0 to −10.0) | −0.53 |
| *Fit quality* |  |  |  |
| Fit *R*<sup>2</sup> ≥ 0.8 | 1,156 | −11.7 (−12.9 to −10.5) | −0.50 |
| Fit *R*<sup>2</sup> ≥ 0.95 | 1,156 | −11.4 (−12.7 to −10.1) | −0.49 |
| Fit *R*<sup>2</sup> ≥ 0.98 | 1,143 | −10.7 (−12.2 to −9.2) | −0.43 |
| Regional mean (not median) of log τ | 1,156 | −11.3 (−12.5 to −10.1) | −0.49 |
| *Covariates* |  |  |  |
| + aperiodic exponent | 1,156 | −8.2 (−9.3 to −7.1) | – |
| + broadband offset | 1,156 | −7.4 (−8.4 to −6.3) | – |
| + alpha peak frequency | 1,155 | −12.1 (−13.3 to −10.9) | – |
| + alpha peak power | 1,155 | −11.1 (−12.3 to −9.9) | – |
| + fit error | 1,156 | −9.4 (−10.6 to −8.1) | – |
| + fit *R*<sup>2</sup> | 1,156 | −6.8 (−8.3 to −5.3) | – |
| + number of valid regions | 1,156 | −11.0 (−12.2 to −9.9) | – |
| + all spectral covariates | 1,155 | −6.2 (−7.2 to −5.1) | – |
| *Site* |  |  |  |
| Cam-CAN only | 606 | −8.9 (−10.5 to −7.3) | −0.43 |
| OMEGA only | 162 | −9.5 (−14.4 to −4.4) | −0.33 |
| PANS only | 388 | −15.5 (−17.5 to −13.4) | −0.64 |
| Leave out Cam-CAN | 550 | −14.4 (−16.2 to −12.4) | −0.56 |
| Leave out OMEGA | 994 | −11.9 (−13.2 to −10.7) | −0.52 |
| Leave out PANS | 768 | −9.0 (−10.5 to −7.4) | −0.41 |
| ComBat-harmonised (age, sex protected) + site term | 1,057 | −11.4 (−12.7 to −10.1) | −0.50 |
| *Statistical model* |  |  |  |
| Robust (Huber) regression | 1,156 | −11.8 (−12.9 to −10.7) | −0.50 |
| *Age range* |  |  |  |
| Adults only (≥ 18 y) | 924 | −8.1 (−9.6 to −6.6) | −0.38 |
| Overlapping range of all sites (18–74 y) | 805 | −9.3 (−11.1 to −7.4) | −0.38 |
| Age ≥ 40 y | 572 | −4.4 (−7.0 to −1.9) | −0.17 |
| Age < 40 y | 584 | −28.0 (−32.2 to −23.4) | −0.41 |
| *Sex* |  |  |  |
| Females only | 562 | −11.3 (−13.2 to −9.4) | −0.47 |
| Males only | 594 | −12.2 (−13.8 to −10.6) | −0.53 |
| *Spectral parameterisation* |  |  |  |
| Fit range 1–45 Hz | 1,154 | −13.0 (−14.5 to −11.4) | −0.44 |
| Fit range 3–40 Hz | 1,156 | −11.2 (−12.4 to −10.1) | −0.53 |
| ≤ 3 peaks, width 2–8 Hz | 1,155 | −11.9 (−13.1 to −10.7) | −0.51 |

*Note. n*, number of participants. % per decade, estimated change in global τ per decade of age. Meta ρ, within-cohort meta-analytic Spearman correlation between age and global τ (Fisher-z, inverse-variance weights); –, not computed. Rows are grouped by type of analysis.

**Supplementary Table 3a.** | Correlations with age for Supplementary Fig. 4.

| Cohort | *n* | Spearman |  |  | Pearson |  |  |
|---|---|---|---|---|---|---|---|
|  |  | ρ | 95% CI | *P* | *r* | 95% CI | *P* |
| *log<sub>10</sub> τ* |  |  |  |  |  |  |  |
| Cam-CAN | 606 | −0.434 | −0.496 to −0.367 | 3.436 × 10<sup>−29</sup> | −0.397 | −0.462 to −0.328 | 2.563 × 10<sup>−24</sup> |
| OMEGA | 162 | −0.333 | −0.464 to −0.189 | 1.464 × 10<sup>−5</sup> | −0.289 | −0.424 to −0.141 | 1.928 × 10<sup>−4</sup> |
| PANS | 388 | −0.640 | −0.695 to −0.577 | 4.875 × 10<sup>−46</sup> | −0.577 | −0.640 to −0.507 | 7.837 × 10<sup>−36</sup> |
| Pooled | 1,156 | −0.654 | −0.686 to −0.619 | 7.525 × 10<sup>−142</sup> | −0.609 | −0.644 to −0.572 | 2.510 × 10<sup>−118</sup> |
| Within-cohort meta-analysis | 1,156 | −0.498 | −0.540 to −0.453 | 1.716 × 10<sup>−76</sup> | – | – | – |
| *Aperiodic exponent* |  |  |  |  |  |  |  |
| Cam-CAN | 606 | 0.364 | 0.293 to 0.432 | 1.785 × 10<sup>−20</sup> | 0.342 | 0.269 to 0.410 | 4.762 × 10<sup>−18</sup> |
| OMEGA | 162 | 0.250 | 0.100 to 0.389 | 0.001 | 0.341 | 0.197 to 0.471 | 8.919 × 10<sup>−6</sup> |
| PANS | 388 | −0.113 | −0.210 to −0.014 | 0.026 | −0.001 | −0.100 to 0.099 | 0.991 |
| Pooled | 1,156 | 0.558 | 0.517 to 0.596 | 1.660 × 10<sup>−95</sup> | 0.538 | 0.496 to 0.578 | 9.880 × 10<sup>−88</sup> |
| Within-cohort meta-analysis | 1,156 | 0.196 | 0.139 to 0.251 | 1.911 × 10<sup>−11</sup> | – | – | – |
| *Alpha peak frequency* |  |  |  |  |  |  |  |
| Cam-CAN | 606 | −0.358 | −0.425 to −0.286 | 9.768 × 10<sup>−20</sup> | −0.361 | −0.428 to −0.289 | 4.726 × 10<sup>−20</sup> |
| OMEGA | 162 | −0.112 | −0.261 to 0.043 | 0.157 | −0.186 | −0.331 to −0.033 | 0.018 |
| PANS | 387 | 0.132 | 0.033 to 0.229 | 0.009 | 0.065 | −0.035 to 0.164 | 0.203 |
| Pooled | 1,155 | −0.214 | −0.268 to −0.158 | 2.083 × 10<sup>−13</sup> | −0.225 | −0.279 to −0.169 | 1.083 × 10<sup>−14</sup> |
| Within-cohort meta-analysis | 1,155 | −0.166 | −0.222 to −0.110 | 1.311 × 10<sup>−8</sup> | – | – | – |

*Note.* Within-cohort meta-analysis, Fisher-z inverse-variance pooling of the three cohort estimates; Pooled, all participants combined, ignoring cohort. –, not applicable.

**Supplementary Table 3b.** | Linear and non-linear age models for Supplementary Fig. 4.

| Cohort | Slope per decade | 95% CI | *P* (slope) | *R*<sup>2</sup> linear | *R*<sup>2</sup> spline | *F* (non-linearity) | *P* (non-linearity) | *I*<sup>2</sup> |
|---|---|---|---|---|---|---|---|---|
| *log<sub>10</sub> τ* |  |  |  |  |  |  |  |  |
| Cam-CAN | −0.041 | −0.048 to −0.033 | 3.020 × 10<sup>−25</sup> | 0.158 | 0.198 | 15.030 | 4.271 × 10<sup>−7</sup> | – |
| OMEGA | −0.040 | −0.064 to −0.016 | 0.001 | 0.083 | 0.093 | 0.793 | 0.454 | – |
| PANS | −0.073 | −0.083 to −0.062 | 2.057 × 10<sup>−42</sup> | 0.333 | 0.407 | 23.931 | 1.604 × 10<sup>−10</sup> | – |
| Pooled | −0.054 | −0.060 to −0.048 | 3.034 × 10<sup>−68</sup> | 0.406 | 0.455 | 51.869 | 2.706 × 10<sup>−22</sup> | – |
| Within-cohort meta-analysis | – | – | – | – | – | – | – | 0.927 |
| *Aperiodic exponent* |  |  |  |  |  |  |  |  |
| Cam-CAN | 0.134 | 0.106 to 0.161 | 1.324 × 10<sup>−21</sup> | 0.117 | 0.143 | 9.357 | 9.956 × 10<sup>−5</sup> | – |
| OMEGA | 0.152 | 0.068 to 0.236 | 4.087 × 10<sup>−4</sup> | 0.116 | 0.139 | 2.072 | 0.129 | – |
| PANS | 0.000 | −0.017 to 0.017 | 0.994 | 0.000 | 0.038 | 7.518 | 6.271 × 10<sup>−4</sup> | – |
| Pooled | 0.080 | 0.062 to 0.098 | 2.684 × 10<sup>−18</sup> | 0.571 | 0.587 | 23.218 | 1.302 × 10<sup>−10</sup> | – |
| Within-cohort meta-analysis | – | – | – | – | – | – | – | 0.966 |
| *Alpha peak frequency* |  |  |  |  |  |  |  |  |
| Cam-CAN | −0.208 | −0.254 to −0.161 | 1.444 × 10<sup>−18</sup> | 0.130 | 0.130 | 0.092 | 0.912 | – |
| OMEGA | −0.150 | −0.276 to −0.025 | 0.019 | 0.035 | 0.069 | 2.883 | 0.059 | – |
| PANS | 0.030 | −0.031 to 0.091 | 0.341 | 0.004 | 0.039 | 6.902 | 0.001 | – |
| Pooled | −0.104 | −0.141 to −0.067 | 3.808 × 10<sup>−8</sup> | 0.091 | 0.135 | 29.577 | 2.983 × 10<sup>−13</sup> | – |
| Within-cohort meta-analysis | – | – | – | – | – | – | – | 0.967 |

*Note.* Slope, change per decade of age (log<sub>10</sub> τ, aperiodic exponent or Hz). *R*<sup>2</sup> linear and *R*<sup>2</sup> spline, variance explained by a linear or a spline age model; *F* and *P* (non-linearity), test of non-linearity (spline versus linear model); *I*<sup>2</sup>, heterogeneity across cohorts in the within-cohort meta-analysis (fraction). –, not applicable.

**Supplementary Table 4.** | Test–retest reproducibility of corticothalamic fits (60 participants re-fitted with an independent library).

| Quantity | Spearman ρ | Pearson *r* |
|---|---|---|
| G<sub>ee</sub> | 0.52 | 0.49 |
| G<sub>ei</sub> | 0.43 | 0.50 |
| G<sub>ese</sub> | 0.85 | 0.84 |
| G<sub>esre</sub> | 1.00 | 0.98 |
| G<sub>srs</sub> | 0.98 | 0.99 |
| α | 0.96 | 0.99 |
| β | 0.33 | 0.72 |
| t<sub>0</sub> | 0.99 | 0.99 |
| EMG<sub>a</sub> | 0.76 | 0.48 |
| X | 0.98 | 1.00 |
| Y | 1.00 | 1.00 |
| Z | 0.99 | 0.99 |

*Note.* Derived quantities: *X* = G<sub>ee</sub>/(1 − G<sub>ei</sub>); *Y* = (G<sub>ese</sub> + G<sub>esre</sub>)/[(1 − G<sub>srs</sub>)(1 − G<sub>ei</sub>)]; *Z* = −G<sub>srs</sub>αβ/(α + β)<sup>2</sup>.

**Supplementary Table 5.** | Age effects on corticothalamic quantities (participant-level fits).

| Quantity | Meta ρ (95% CI) | *q* | Spearman ρ by cohort |  |  |
|---|---|---|---|---|---|
|  |  |  | Cam-CAN | OMEGA | PANS |
| X | −0.11 (−0.17 to −0.05) | <0.001 | −0.037 | −0.057 | −0.239 |
| Y | 0.15 (0.09 to 0.21) | <0.001 | 0.132 | −0.147 | 0.293 |
| Z | 0.42 (0.37 to 0.47) | <0.001 | 0.271 | 0.309 | 0.652 |
| G<sub>srs</sub> | −0.48 (−0.52 to −0.43) | <0.001 | −0.328 | −0.373 | −0.689 |
| α | −0.33 (−0.38 to −0.28) | <0.001 | −0.284 | −0.215 | −0.450 |
| β | −0.01 (−0.07 to 0.05) | 0.774 | −0.018 | 0.176 | −0.070 |
| t<sub>0</sub> | 0.09 (0.03 to 0.15) | 0.004 | 0.127 | 0.142 | 0.005 |
| EMG<sub>a</sub> | −0.03 (−0.09 to 0.03) | 0.322 | −0.073 | −0.061 | 0.042 |
| G<sub>ee</sub> | −0.03 (−0.09 to 0.02) | 0.322 | −0.031 | −0.077 | −0.022 |
| G<sub>ei</sub> | −0.02 (−0.08 to 0.04) | 0.599 | 0.014 | 0.011 | −0.080 |
| G<sub>ese</sub> | 0.20 (0.14 to 0.26) | <0.001 | 0.243 | 0.028 | 0.204 |
| G<sub>esre</sub> | 0.16 (0.10 to 0.21) | <0.001 | 0.122 | 0.124 | 0.230 |

*Note.* Meta ρ, within-cohort meta-analytic Spearman correlation with age (Fisher-z, inverse-variance weights); *q*, Benjamini–Hochberg FDR-adjusted *P* value of the meta-analytic estimate. Derived quantities as in Supplementary Table 4.

**Supplementary Table 6.** | In-silico perturbations of the fitted corticothalamic model.

| Shift (age 25 → 75) | Model τ (ms) | Alpha frequency (Hz) | Δτ (%) | Δ alpha frequency (Hz) |
|---|---|---|---|---|
| Young-adult median (20–29 y) | 14.05 | 9.44 | 0.00 | 0.00 |
| G<sub>ee</sub> | 13.88 | 9.45 | −1.22 | 0.01 |
| G<sub>ei</sub> | 13.81 | 9.48 | −1.74 | 0.04 |
| G<sub>ese</sub> | 13.37 | 9.47 | −4.81 | 0.03 |
| G<sub>esre</sub> | 14.05 | 9.44 | 0.00 | 0.00 |
| G<sub>srs</sub> | 9.05 | 9.47 | −35.56 | 0.03 |
| α | 15.71 | 9.65 | 11.83 | 0.21 |
| β | 14.05 | 9.44 | 0.00 | 0.00 |
| t<sub>0</sub> | 13.88 | 9.26 | −1.21 | −0.18 |
| EMG<sub>a</sub> | 14.27 | 9.45 | 1.57 | 0.01 |
| All except G<sub>srs</sub> | 14.64 | 9.41 | 4.23 | −0.03 |
| G<sub>srs</sub> + t<sub>0</sub> only | 8.86 | 9.31 | −36.96 | −0.13 |
| All parameters shifted | 8.86 | 9.61 | −36.96 | 0.17 |
| Observed age 25 → 75 (τ) | – | – | −47.06 | – |
| Observed age 25 → 75 (alpha frequency) | – | – | – | −0.50 |
| Fitted spectra, 20–29 → 70–89 y (full model, group medians) | – | – | −38.40 | −0.73 |
| MEG spectra, 20–29 → 70–89 y (group medians) | – | – | −44.73 | −1.31 |

*Note.* Model τ and alpha frequency were obtained from the corticothalamic model with the indicated parameter(s) shifted from age 25 to age 75 y (baseline, the median of the nine fitted parameters at 20–29 y; shift, age slope × 50 y, clipped to the parameter bounds); Δ, change relative to the young-adult median. Parameters are shifted independently, so correlations among parameters are ignored; β and G<sub>esre</sub> are at a bound in most fits and therefore change little. “Observed” rows give the empirical change between 25 and 75 y from the fitted age trajectory. The last two rows compare the age change in τ and alpha frequency of the fitted model spectra with that of the MEG spectra directly (group medians pooled over cohorts, n = 188 and 180). –, not applicable.

**Supplementary Table 7.** | Ageing knock-in analyses in the 68-module RNN.

| Aged parameter(s) | Median Δτ (%/decade) | Regions shorter (of 68) | Empirical age map |  | Gradient |  | Visual (%/decade) | Other (%/decade) |
|---|---|---|---|---|---|---|---|---|
|  |  |  | ρ | *P*<sub>spin</sub> | ρ | *P*<sub>spin</sub> |  |  |
| All parameters (old fits) | −8.48 | 66 | 0.72 | 0.02 | −0.52 | 0.01 | −0.44 | −9.15 |
| G<sub>ee</sub> | 3.40 | 28 | unstable | – | unstable | – | 2.65 | 3.40 |
| G<sub>ei</sub> | −0.48 | 36 | unstable | – | unstable | – | 0.09 | −1.13 |
| G<sub>ese</sub> | −1.66 | 47 | 0.59 | <0.01 | 0.01 | 0.97 | −0.06 | −1.80 |
| G<sub>esre</sub> | 0.00 | 42 | −0.26 | 0.13 | 0.23 | 0.12 | 0.00 | 0.00 |
| G<sub>srs</sub> | −3.03 | 56 | −0.12 | 0.76 | 0.26 | 0.24 | −1.35 | −4.45 |
| α | 0.04 | 29 | 0.39 | 0.30 | −0.14 | 0.54 | 0.60 | 0.00 |
| β | 0.00 | 45 | −0.07 | 0.72 | −0.04 | 0.78 | 0.00 | 0.00 |
| t<sub>0</sub> | −0.52 | 46 | −0.41 | 0.13 | −0.08 | 0.73 | −0.52 | −0.43 |

*Note.* Regions shorter, number of the 68 regions in which τ was shorter after ageing. unstable, network unstable under the perturbation (ρ and *P*<sub>spin</sub> not computed; –). *P*<sub>spin</sub>, spin-test *P* value.

**Supplementary Table 8.** | Reservoir analyses of the connectome thalamocortical RNN.

| Network | Spontaneous knee τ (ms) | Memory capacity (mean ± s.d.) | Δ capacity vs young (95% CI) | Integration *R*<sup>2</sup> (500 ms) |
|---|---|---|---|---|
| G<sub>srs</sub> × 0.25 | 36.7 | 3.57 ± 0.17 | −0.94 (−0.99 to −0.88) | 0.762 |
| G<sub>srs</sub> × 0.5 | 27.3 | 4.03 ± 0.18 | −0.48 (−0.52 to −0.44) | 0.778 |
| G<sub>srs</sub> × 1.5 | 10.5 | 4.85 ± 0.14 | +0.34 (+0.31 to +0.37) | 0.798 |
| G<sub>srs</sub> × 2.0 | 6.1 | 5.13 ± 0.13 | +0.62 (+0.57 to +0.67) | 0.804 |
| Old | 11.6 | 4.64 ± 0.18 | +0.13 (+0.09 to +0.17) | 0.793 |
| Young | 16.0 | 4.51 ± 0.16 | – | 0.777 |
| Young + aged G<sub>srs</sub> | 10.0 | 4.83 ± 0.16 | +0.33 (+0.29 to +0.36) | 0.797 |
| Young + aged unit time constants | 17.5 | 4.49 ± 0.17 | −0.02 (−0.03 to −0.00) | 0.775 |

*Note.* G<sub>srs</sub> × *k*, intrathalamic loop gain scaled by a factor *k* in the young network. Δ capacity, difference in memory capacity from the young network (95% CI).

**Supplementary Table 9.** | Nested comparison of cortex-only and corticothalamic neural-field models.

| Model | *k* | In-sample *R*<sup>2</sup>, median (IQR) | ΔBIC vs M3, median | Lowest BIC (%) | Held-out RMSE, median (IQR) | Alpha peak (%) | Valid τ (%) | ρ, model vs MEG τ | ρ, age vs τ |
|---|---|---|---|---|---|---|---|---|---|
| M0 Cortex only | 5 | 0.691 (0.512–0.801) | 122 | 0.1 | 0.328 (0.257–0.410) | 15 | 15 | −0.15 (−0.30 to −0.01) | 0.13 (−0.02 to 0.27) |
| M1 + G<sub>ese</sub>, t<sub>0</sub> | 7 | 0.758 (0.626–0.845) | 119 | 0.0 | 0.309 (0.246–0.379) | 77 | 20 | −0.11 (−0.24 to 0.02) | 0.02 (−0.12 to 0.15) |
| M2 + G<sub>esre</sub> | 8 | 0.745 (0.606–0.836) | 124 | 0.0 | 0.315 (0.252–0.389) | 77 | 13 | −0.23 (−0.37 to −0.07) | 0.09 (−0.07 to 0.25) |
| M3 Full (+ G<sub>srs</sub>) | 9 | 0.985 (0.974–0.992) | reference | 99.9 | 0.141 (0.092–0.220) | 91 | 89 | 0.87 (0.86 to 0.88) | −0.33 (−0.38 to −0.27) |
| MEG spectra | – | – | – | – | – | 92 | 94 | – | −0.42 (−0.47 to −0.37) |

*Note. k*, number of free parameters. All models were fitted to the cortex-averaged spectrum (1–45 Hz, 45 bins) of every participant (n = 1,160; 1,156 with age plus four OMEGA participants without age). Held-out RMSE (log<sub>10</sub> power) is from blocked cross-validation over five 9-Hz blocks in 300 participants (100 per cohort); IQR, interquartile range. ΔBIC uses n = 45 bins and is descriptive because adjacent bins are correlated.
*Note.* Alpha peak and valid τ give the percentage of spectra in which a 7–14 Hz peak, or a knee with τ < 500 ms and R<sup>2</sup> ≥ 0.9, could be recovered by spectral parameterization (the same procedure as for MEG). ρ is the Spearman correlation pooled within cohort by inverse-variance weighting on the Fisher z scale (95% confidence interval), computed over participants with valid estimates in both spectra (model vs MEG τ) or for age (age vs τ).

## Notes

https://github.com/Laoma29/MEG_Timescale_networ

### Competing Interest Statement

The authors have declared no competing interest.

## References

1. Murray, J. D. et al. A hierarchy of intrinsic timescales across primate cortex. Nat. Neurosci. 17, 1661–1663 (2014).

2. Gao, R., van den Brink, R. L., Pfeffer, T. & Voytek, B. Neuronal timescales are functionally dynamic and shaped by cortical microarchitecture. eLife 9, e61277 (2020).

3. Zeraati, R., Levina, A., Macke, J. H. & Gao, R. Neural timescales from a computational perspective. Nat. Neurosci. 29, 1534–1547 (2026).

4. Rempe, M. P., et al. Spontaneous cortical dynamics from the first years to the golden years. Proc. Natl Acad. Sci. USA 120, e2212776120 (2023).

5. Chaudhuri, R., Knoblauch, K., Gariel, M.-A., Kennedy, H. & Wang, X.-J. A large-scale circuit mechanism for hierarchical dynamical processing in the primate cortex. Neuron 88, 419–431 (2015).

6. Wang, X.-J. Macroscopic gradients of synaptic excitation and inhibition in the neocortex. Nat. Rev. Neurosci. 21, 169–178 (2020).

7. Steriade, M., McCormick, D. A. & Sejnowski, T. J. Thalamocortical oscillations in the sleeping and aroused brain. Science 262, 679–685 (1993).

8. Sherman, S. M. & Guillery, R. W. The role of the thalamus in the flow of information to the cortex. Phil. Trans. R. Soc. Lond. B 357, 1695–1708 (2002).

9. Halassa, M. M. & Sherman, S. M. Thalamocortical circuit motifs: a general framework. Neuron 103, 762–770 (2019).

10. Guo, Z. V. et al. Maintenance of persistent activity in a frontal thalamocortical loop. Nature 545, 181–186 (2017).

11. Schmitt, L. I. et al. Thalamic amplification of cortical connectivity sustains attentional control. Nature 545, 219–223 (2017).

12. Bolkan, S. S. et al. Thalamic projections sustain prefrontal activity during working memory maintenance. Nat. Neurosci. 20, 987–996 (2017).

13. Hughes, E. J. et al. Regional changes in thalamic shape and volume with increasing age. NeuroImage 63, 1134–1142 (2012).

14. Fama, R. & Sullivan, E. V. Thalamic structures and associated cognitive functions: relations with age and aging. Neurosci. Biobehav. Rev. 54, 29–37 (2015).

15. Robinson, P. A. et al. Prediction of electroencephalographic spectra from neurophysiology. *Phys*. Rev. E 63, 021903 (2001).

16. Robinson, P. A., Rennie, C. J., Rowe, D. L. & O’Connor, S. C. Estimation of multiscale neurophysiologic parameters by electroencephalographic means. Hum. Brain Mapp. 23, 53–72 (2004).

17. Abeysuriya, R. G. & Robinson, P. A. Real-time automated EEG tracking of brain states using neural field theory. J. Neurosci. Methods 258, 28–45 (2016).

18. van Albada, S. J., Kerr, C. C., Chiang, A. K. I., Rennie, C. J. & Robinson, P. A. Neurophysiological changes with age probed by inverse modeling of EEG spectra. Clin. Neurophysiol. 121, 21–38 (2010).

19. Halgren, M., et al. The generation and propagation of the human alpha rhythm. Proc. Natl Acad. Sci. USA 116, 23772–23782 (2019).

20. Shafto, M. A. et al. The Cambridge Centre for Ageing and Neuroscience (Cam-CAN) study protocol: a cross-sectional, lifespan, multidisciplinary examination of healthy cognitive ageing. BMC Neurol. 14, 204 (2014).

21. Taylor, J. R. et al. The Cambridge Centre for Ageing and Neuroscience (Cam-CAN) data repository: structural and functional MRI, MEG, and cognitive data from a cross-sectional adult lifespan sample. NeuroImage 144, 262–269 (2017).

22. Niso, G. et al. OMEGA: the Open MEG Archive. NeuroImage 124, 1182–1187 (2016).

23. Tadel, F., Baillet, S., Mosher, J. C., Pantazis, D. & Leahy, R. M. Brainstorm: a user-friendly application for MEG/EEG analysis. Comput. Intell. Neurosci. 2011, 879716 (2011).

24. Gross, J. et al. Good practice for conducting and reporting MEG research. NeuroImage 65, 349–363 (2013).

25. Fischl, B. FreeSurfer. NeuroImage 62, 774–781 (2012).

26. Donoghue, T. et al. Parameterizing neural power spectra into periodic and aperiodic components. Nat. Neurosci. 23, 1655–1665 (2020).

27. Margulies, D. S., et al. Situating the default-mode network along a principal gradient of macroscale cortical organization. Proc. Natl Acad. Sci. USA 113, 12574–12579 (2016).

28. Van Essen, D. C. et al. The WU-Minn Human Connectome Project: an overview. NeuroImage 80, 62–79 (2013).

29. Larivière, S. et al. The ENIGMA Toolbox: multiscale neural contextualization of multisite neuroimaging datasets. Nat. Methods 18, 698–700 (2021).

30. Benjamini, Y. & Hochberg, Y. Controlling the false discovery rate: a practical and powerful approach to multiple testing. J. R. Stat. Soc. B 57, 289–300 (1995).

31. Alexander-Bloch, A. F. et al. On testing for spatial correspondence between maps of human brain structure and function. NeuroImage 178, 540–551 (2018).

32. Johnson, W. E., Li, C. & Rabinovic, A. Adjusting batch effects in microarray expression data using empirical Bayes methods. Biostatistics 8, 118–127 (2007).

33. Fortin, J.-P. et al. Harmonization of cortical thickness measurements across scanners and sites. NeuroImage 167, 104–120 (2018).

34. Raut, R. V., Snyder, A. Z. & Raichle, M. E. Hierarchical dynamics as a macroscopic organizing principle of the human brain. Proc. Natl Acad. Sci. USA 117, 20890–20897 (2020).

35. Demirtaş, M. et al. Hierarchical heterogeneity across human cortex shapes large-scale neural dynamics. Neuron 101, 1181–1194.e13 (2019).

36. Honey, C. J., Kötter, R., Breakspear, M. & Sporns, O. Network structure of cerebral cortex shapes functional connectivity on multiple time scales. Proc. Natl Acad. Sci. USA 104, 10240–10245 (2007).

37. Voytek, B. et al. Age-related changes in 1/f neural electrophysiological noise. J. Neurosci. 35, 13257–13265 (2015).

